# Ki-67 is necessary during DNA replication for forks protection and genome stability

**DOI:** 10.1101/2023.04.18.537310

**Authors:** Konstantinos Stamatiou, Florentin Huguet, Christos Spanos, Juri Rappsilber, Paola Vagnarelli

## Abstract

**Background:** The proliferation antigen Ki-67 has been widely used in clinical settings for cancer staging for many years but investigations on its biological functions have lagged. Recently, Ki-67 was shown to regulate both the composition of the chromosome periphery and chromosome behaviour in mitosis as well as to play a role in heterochromatin organisation and gene transcription. However, a role for Ki-67 in regulating cell cycle progression has never been reported. The progress towards understanding Ki-67 function have been limited by the tools available to deplete the protein coupled to its abundance and fluctuation during the cell cycle.

**Results:** Here we have used an auxin-inducible degron tag (AID) to achieve a rapid and homogeneous degradation of Ki-67 in HCT116 cells. This system, coupled with APEX2 proteomics and phospho-proteomics approaches, allowed us to show for the first time that Ki-67 plays a role during DNA replication. In its absence, DNA replication is severely delayed, the replication machinery is unloaded, causing DNA damage that is not sensed by the canonical pathways and dependant on HUWE1 ligase. This leads to replication and sister chromatids cohesion defects, but it also triggers an interferon response mediated by the cGAS/STING pathway in all the cell lines tested.

**Conclusions:** We have unveiled a new function of Ki-67 in DNA replication and genome maintenance that is independent of its previously known role in mitosis and gene regulation.

## Background

Since its discovery in 1983 [1], Ki-67 has been widely used as proliferation marker and adopted by pathologists and clinicians to stage multiple cancer types [2–4] [5] [6, 7] but the role for this protein in cell cycle regulation has yet to be found. Although it is now well accepted that Ki-67 is important for the organisation of the chromosome periphery in mitosis [8] [9, 10] and its surfactant properties allow the chromosomes to both maintain separation during early mitosis [11] and coalesce during mitotic exit [12], cells seem to proliferate almost normally in the absence of this marker [11] [13]. The ability of cells to progress in the cell cycle without Ki-67 has been a matter of controversy for some years (for reviews discussing this topic see [14–17]). In summary, it appears that oligo-based approaches for depleting Ki-67 lead to a proliferative disadvantage in cancer cells [18] while Ki-67 knock-out cell lines are viable in human [11] and mouse [13]. However, these KO cells behave differently and present broad changes in hundreds of transcripts [13], they cannot metastasise in orthotopic cancer models [19, 20] while mice without Ki-67 are more resistant to cancer development [19, 21–23]. The mechanism explaining these phenotypes is still obscure, although it could be extremely valuable in terms of understanding the role of Ki-67 in cancer or exploiting this marker for therapy. Moreover, since Ki-67 expression predicts the differential response of cell lines to CDK inhibitors treatment during xenograft tumour formation [24], understanding how Ki-67 expression and sensitivity to its depletion are linked, is an important goal for developing stratified approaches to cancer therapies.

Ki-67 levels are regulated during the cell cycle. During mitotic exit and early G1 the protein is degraded via the ubiquitin-proteosome system [24–26] and, upon passage through the G1 restriction point, CDK4/6 activation triggers Ki-67 transcription. Therefore, it seems that its levels are indeed linked to the G1/S cell cycle progression but no role for Ki-67 has been so far demonstrated in this cell cycle transition.

Another phenotype linked to Ki-67 depletion is a compromised heterochromatin maintenance as revealed by the Ki-67 KO mouse model [13] or the RNAi study in hTERT-RPE1 cells [27] and positioning [28].

The link between all the different phenotypes observed in Ki-67 depletion or Knock-out models is currently not very clear. The key outstanding question is: are those phenotypes dependent on Ki-67 function at the chromosome periphery and linked to the re-establishment of chromatin organisation in G1 or is the protein playing several roles during the cell cycle that lead to different outcomes depending on the genetic background and intrinsic compensatory effects?

RNAi-based depletions cannot answer the question and, similarly, the selection for viability could lead to compensatory mechanisms.

Here, we have addressed the question of understanding the role of Ki-67 at the G1/S transition and DNA replication using a newly generated endogenously degron tagged cell line for Ki-67 where rapidly (in 4 h), and homogenously, we can deplete Ki-67 at the G1/S boundary, thus separating its effect from possible other functions in other cell cycle stages. Using this system, we discovered a novel function of Ki-67 during DNA replication. Being also endogenously tagged with GFP, we could show that Ki-67 localisation changes concomitantly with late replicating regions. Here, we provide evidence that degradation of Ki-67 at the G1/S boundary delays replication and causes DNA damage. The replication forks are unprotected, and DNA replication is incomplete with defects in cohesion maintenance. These effects are also separated from the documented Ki-67 function on transcription. However, the delay in cell cycle progression is overcome and, eventually, cells start growing albeit this triggers a persistent interferon response mediated by the cGAS/STING pathway.

## RESULTS

### Ki-67 is important for timely progression of DNA replication

Ki-67 has been linked to chromatin organisation and transcription regulation in many systems [13, 27]. However, the most investigated aspect of Ki-67 biology has been its role at the chromosome periphery during mitosis [8] [11] [16] [9] [15]. Most experiments involved siRNA or ShRNA mediated depletion over a long period of time or KO cell lines (which have been selected for and could have led to adaptation) [11] [13]. More recently, the use of a degron system has allowed to dissect more specifically some of the functions of Ki-67 in mitosis [10], but the cell line available to the community, although extremely valuable for single cell studies, appears to have a varied response to auxin where some cells are not responding [10], making any biochemical approach difficult to interpret.

To overcome this problem, we have generated another endogenously-tagged:AID cell line in HCT116 where both Ki-67 alleles are fused to the mClover:AID module. In this cell line, OSTR is expressed under a doxycycline (Dox) inducible promoter; addition of Dox and Auxin leads to Ki-67 degradation (Figure 1 A, and Supplementary Figure 1 A, B). After several rounds of subcloning, we isolated cell lines where Ki-67 degradation occurs homogenously within 3 h (Figure 1 B, and Supplementary Figure 1 C) after auxin treatment. From now on, any reference to auxin treatment refers to the addition of both dox and auxin to the culture. Using this cell line (Ki-67-AID), we first assessed the ability of cells to survive when Ki-67 degradation is maintained for several days. In these conditions, the proliferation of cells lacking Ki-67 is severely impaired up to 96 h (Figure 1 C); however, we noticed an increase in cell number at 120 h, suggesting that cells might adapt to the lack of Ki-67 over time. Interestingly, we were not able to obtain Ki-67-AID tagged clones in the OSTR constitutively expressing cell line, further suggesting an important role of Ki-67 for cell survival. Cell cycle analyses by flow cytometry did not show sign of apoptosis but clearly revealed that the proportion of cells in S phase was greatly diminished (Figure 1 D). This new cell line therefore offered us the opportunity to investigate the role of Ki-67 in cell cycle transitions. Being Ki-67 the reference standard for proliferation markers, where its absence is associated with quiescence or senescence [25, 26], and the observed reduced S phase population upon its degradation, we focused on its role at the G1/S transition.

**Figure 1.**
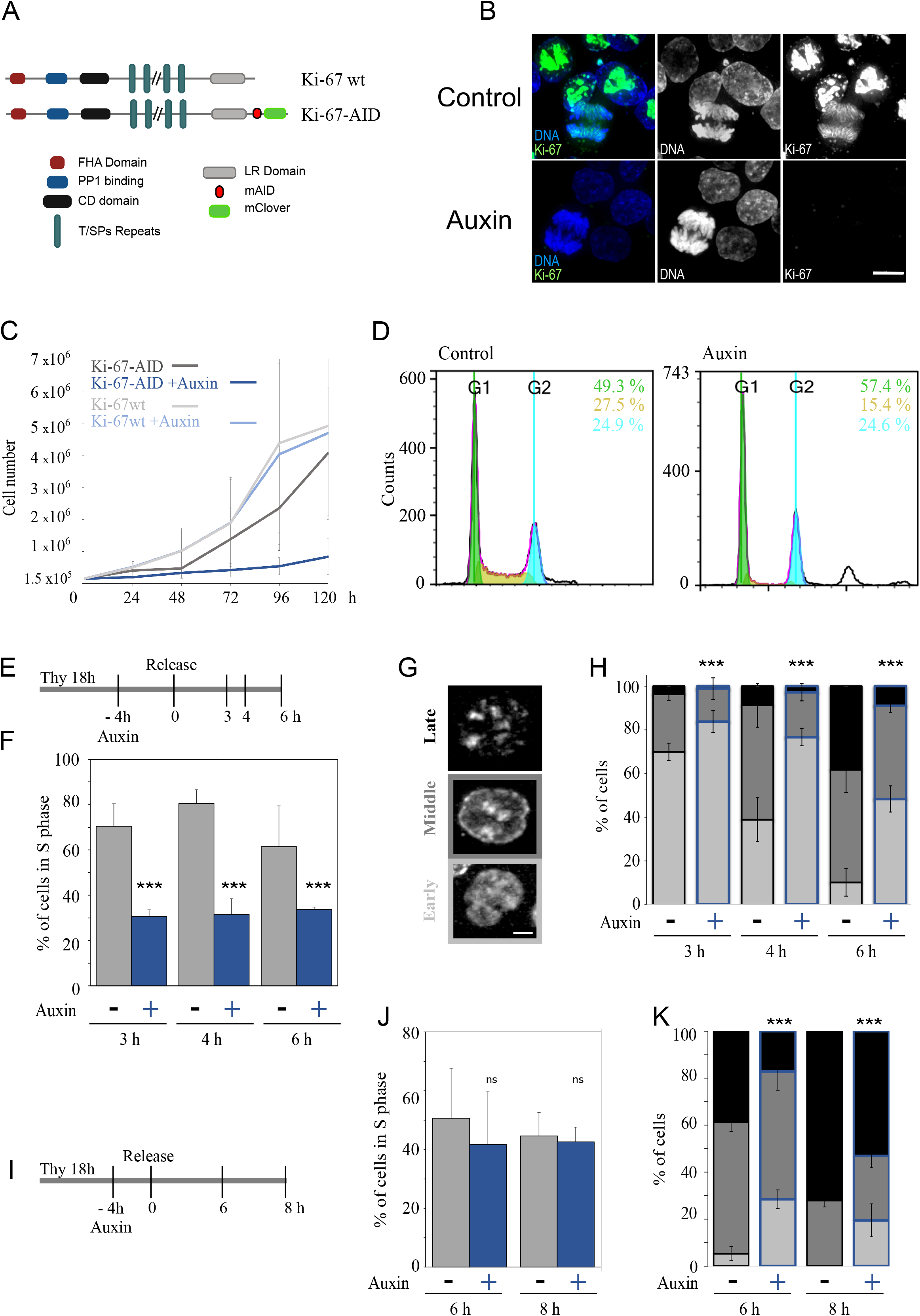
Ki-67 degradation delays S phase progression. A) Scheme of Ki-67 protein structure and domains (Ki-67 wt) and the endogenously tagged version (Ki-67-AID) generated in the HCT116 cell line B) Representative image of the HCT116:Ki-67-AID cell line without (top panels) and with (bottom panels) Auxin. Ki-67 is in green and DNA in blue. Scale bar 10μm. C) Growth curves of the HCTT116 parental (Ki-67 wt) and degron tagged HCT116:Ki-67-AID (Ki-67-AID) cell lines without and with (+ Auxin) doxycycline and Auxin. The values represent the mean of 4 independent experiments. The error bars represent the standard deviations. D) Flowcytometry profiles of the HCT116:Ki-67-AID cell line grown with (Auxin – right) and without (control – left) Auxin. The numbers represent the percentage of cells in each stage: green G1; yellow S; cyan G2/M. E) Scheme of the experiment for F-H. EdU was added 30’ before each time point. Thy= thymidine. F) The graph represents the percentage of cells EdU positive at the different time points. The values are the mean of 5 biological replicas and the error bars represent the standard deviations. Sample sizes: Control 3h=723, 4h=922, 6h=847 Auxin 3h=1207, 4h=1108, 6h=1037. The data were statistically analysed with a Chi squared test (control vs auxin). ***= p<0.001. G) Representative images of EdU patterns for early (bottom panel), middle (middle panel) and late (top panel). H) Distribution of the replicating cells according to the patterns shown in (G) from the experiment in F-H. I) Scheme of the experiment for J-K. EdU was added 30’ before each time point. Thy = thymidine. J) The graph represents the percentage of EdU positive cells at the different time points. The values are the mean of 3 biological replicas and the error bars represent the standard deviations. Samples size: Control: 6h=570, 8h=758, Auxin: 6h=945, 8h=813.The data were statistically analysed with a Chi-squared test (control vs auxin). ***= p<0.001. K) Distribution of the replicating cells according to the patterns shown in (G) from the experiment in J-K.

We wanted to evaluate if the S phase decrease was because of an unknown role of Ki-67 in DNA replication, or a consequence of defects originated in mitosis that had a repercussion on S phase progression or on the transition through the G1 restriction point. To address this, we synchronised cells with thymidine, degraded Ki-67 for 4 h before the release and then monitor S phase progression by EdU incorporation (Figure 1 E, F). Upon release form thymidine, cells without Ki-67 were not efficient in resuming DNA replication even 6 h after release. DNA replication occurs in a sequential and organised manner within the nuclear space and early, middle, and late replication patterns can be identified via EdU incorporation and click chemistry (Figure 1 G). Using this method, we have classified the replication pattern distributions in cells with and without Ki-67 in the same conditions as in Figure 1 (E, F). While control cells progressed through replication as expected, Ki-67 depleted cells showed major delays in progressing form the early to the middle patterns (Figure 1 H), indicating the need for Ki-67 for a timely replication resumption after release from thymidine. To understand if the observed delay was caused by the thymidine treatment, we released cells from the block and then degraded Ki-67 during replication (Figure 1 I); in this condition, the total number of EdU positive cells did not change (Figure 1 J) but the progression of replication was still significantly delayed (Figure 1 K). Furthermore, in experiments where the CDK4/6 inhibitor Palbociclib (supplementary Figure 1 D) was used to arrest cells, we obtained the same results, indicating that the effect is not just a consequence of the thymidine treatment. Moreover, the same phenotype was also observed upon Ki-67 depletion by RNAi in HCT116 using two different previously published oligos against Ki-67 (Supplementary Figure 1 E) [9] [11].

We therefore conclude that lack of Ki-67 affects S phase progression, and this effect is not linked to the passage through mitosis.

### Ki-67 degradation leads to unloading of the replication machinery

DNA damage at G1/S delays S phase progression: this could lead to p21 increase and phosphorylation of H2AX on S139 (γH2AX). We therefore checked p21 and γH2AX levels in cells at the G1/S boundary with and without Ki-67. We did not observe p21 or γH2AX increase in cells lacking Ki-67 but, on the contrary, a decrease in both these markers (Figure 2 A). We then investigated the proteins involved in replication. We analysed by quantitative western blot (LICOR) the levels of chromatin-associated ORC1, MCM3 and PCNA to monitor several stages of origin assembly and firing (Figure 2 B) at the G1/S boundary with and without Ki-67. The analyses showed reduced chromatin-bound levels for all the components in cells treated with auxin (Figure 2 C, D). As ORC1 is already loaded when the thymidine block is applied (and before Ki-67 degradation), its decrease upon Ki-67 depletion, suggests that the replication machinery is unloaded after Ki-67 degradation. Because Ki-67 has also been linked to the maintenance of heterochromatin [13, 27], we considered the possibility that Ki-67 degradation could lead to epigenetic changes in the chromatin that ultimately could affect DNA replication maybe by destabilising the origins. However, we tested several epigenetic markers including H3K9me2, H3K9me3, H3K27me2/3 and H4K20me1 but we could not detect any difference between cells with and without Ki-67 (Figure 2 E, F). Therefore, chromatin reorganisation is unlikely to be the cause of the replication block. Moreover, no changes in the nucleoli morphology could be observed upon Ki-67 degradation as assessed by nucleolin staining (Supplementary Figure 1 F).

**Figure 2.**
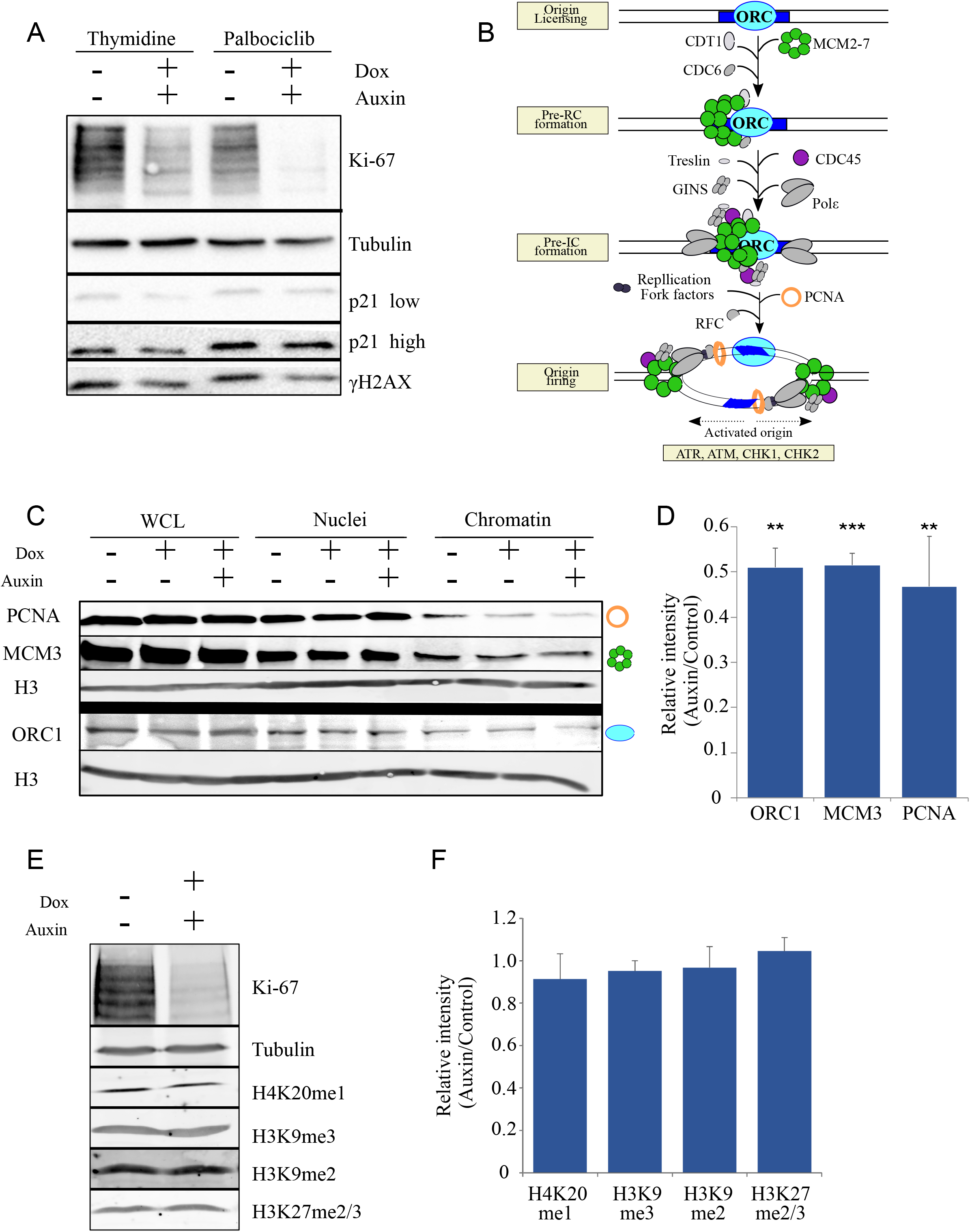
Ki-67 degradation leads to collapse of the replication machinery. A) Western blot of HCT116:Ki-67-AID cell line blocked with either thymidine or Palbociclib 18 h and then treated with or without Auxin for 4 h. The blots were probed with antibodies against Ki-67 (first panel), alpha-tubulin (second panel), p21 (third and fourth panels), and γH2AX (fifth panel). B) Scheme of how the replisome sequentially assembles and activates. C) Representative Western blot of cell fractions (WCL= whole cell lysate, nuclei, and chromatin) of HCT116:Ki-67-AID cell line blocked with thymidine for 18 h and then treated with or without Auxin for 4 h. The blots were probed with antibodies against PCNA (first panel), MCM3 (second panel), Histone H3 (third and fifth panels), and ORC1 (fourth panel). The images were acquired with a LICOR machine in the linear range for quantification purposes. The graphic symbol on the right represents the different proteins depicted in the scheme in (B). D) Quantification of the blot in (C). The values represent the average of 3 independent replicas and the error bars the standard deviations. The experiments were analysed by a Student’s t-test. **= p< 0.01, ***= p< 0.001. E) Representative Western blot of WCL of the HCT116:Ki-67-AID cell line blocked with thymidine 18 h and then treated with or without Auxin for 4 h. The blots were probed with antibodies against Ki-67 (first panel), alpha-tubulin (second panel), H3K20me1 (third panel), H3K9me3 (fourth panel), H3k9me2 (fifth panel) and H3k27me2/3 (sixth panel). The images were acquired with a LICOR machine in the linear range for quantification purposes. F) Quantification of the blot in (E). The values represent the average of 3 independent replicas and the error bars the standard deviations. The experiments were analysed by a Student’s t-test and did not display any statistical significant changes (ns)

These experiments although suggest that Ki-67 is necessary for protecting the replisome and it is important for timely replication progression.

### Ki-67 degradation at the G1/S boundary causes DNA damage and triggers an interferon response

Transcription and replication are coupled and interference with the transcription machinery could directly affect replication or the availability of replication proteins. Because Ki-67 has been linked to transcription regulation [19] [13] [27], we conducted RNA-sequencing experiments in cells at the G1/S boundary with and without Ki-67 or, as a comparison, cells that have passed through mitosis in the absence of Ki-67 (cells synchronised with nocodazole, Ki-67 degradation in nocodazole, released in thymidine). While we observed significant changes in the transcription profile of cells that exited mitosis without Ki-67 (Supplementary Figure 2 A-C), we detected much less changes in transcription for cells where Ki-67 was degraded at the G1/S boundary (Figure 3 A and Supplementary Figure 2 C); in particular, we did not observe downregulation of replication factors suggesting that transcription alterations of essential replication components are not the cause of the problem observed in S phase. However, analyses of the transcripts upregulated in this condition, highlighted genes involved in the viral DNA sensing pathway (Figure 3 B). MAGIC analyses [29] of the regulatory elements present in the genes that change expression upon Ki-67 degradation showed a significant enrichment for STAT1, STAT2, IRF1 and IRF2 for the upregulated genes but no feature was identified for the downregulated ones (Supplementary Figure 2 D). These findings could either represent a specific repressive role of Ki-67 at these genes or indicate the presence of DNA damage that triggers the response. To discriminate between the two possibilities and to validate the findings, we used a reporter plasmid where luciferase expression is driven by the interferon β promoter (Figure 3 C) [30]. Cells were transfected with the plasmid (plus Renilla for normalisation purposes), blocked with thymidine overnight, then auxin was added for 4 h and the cells were released in RO3306 for 18 h before measuring the luciferase induction; in these experimental settings, luciferase expression was triggered in the absence of Ki-67, thus suggesting that Ki-67 degradation activates the interferon response pathway via signalling rather than via a transcriptional repression attenuation. This is the first report of interferon signalling activation caused by lack of Ki-67.

**Figure 3.**
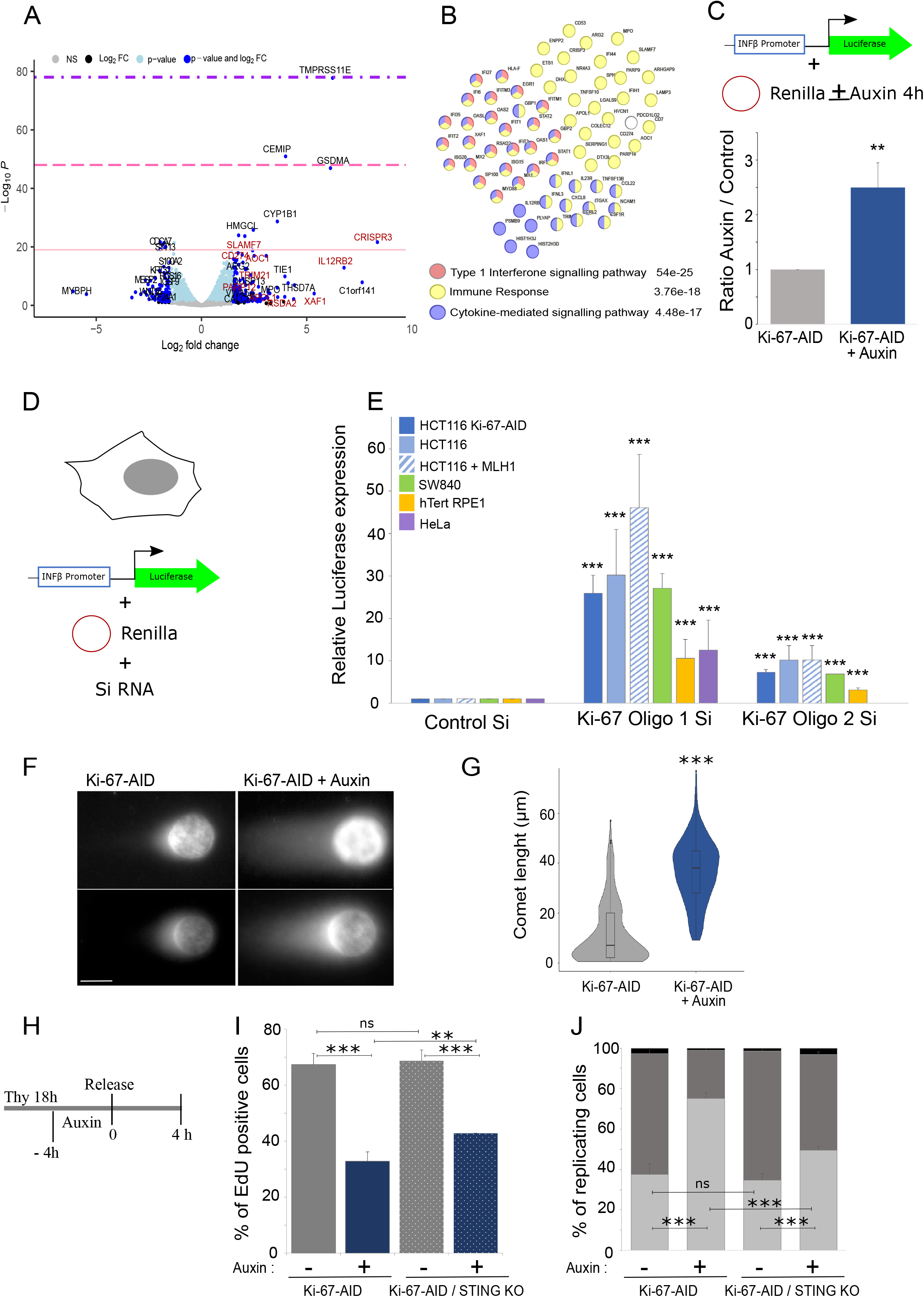
Ki-67 degradation activates the interferon response pathway. A) Volcano plot of the differentially expressed genes obtained by RNA-seq of the HCT116:Ki-67-AID cell line blocked with thymidine for 18 h and then treated with or without Auxin for 4 h. In red are indicated genes that belong to the interferon response. The pink line represents p-value < 10e^-20^, the Hot pink line p-value=10e^-20^-10e^-30^ and the purple line p-value=10e^-20^-10e^-60^. B) STRING analyses of the upregulated genes. The numbers next to the categories represent the false discovery rate. C) HCT116:Ki-67-AID cells were transfected with a plasmid carrying the luciferase gene under the control of the Interferon β promoter together with a plasmid carrying renilla. Cells were treated with thymidine for 18 h, then with auxin for 4 h, and released in RO3306 for 18 h. The graph represents the luciferase activation normalised to renilla in cells untreated (Ki-67-AID) or treated with Auxin for 4 h (Ki-67-AID auxin). The values represent the average of 3 independent experiments and the error bars are the standard deviations. The experiments were analysed by a Student’s t-test. **= p< 0.01. D) Scheme for the transfection set-up used for the experiments in (E). E) Quantification of the luciferase essays performed as in (D) at 72 h post-transfection for the different cell lines. Oligo 1 and Oligo 2 are two independent oligos against Ki-67. The values represent the average of 3 independent replicas and the error bars are the standard deviations. The experiments were analysed by a Student’s t-test. ***= p< 0.001. F) Representative images of the comet essay of the HCT116:Ki-67-AID cell line blocked with thymidine 18 h and then treated with or without Auxin for 4 h. G) Quantification of the comet length. The violin plots represent the distribution of the comet tail length in μm. The box inside the violin represents the 75^th^ and 25^th^ percentile, whiskers are the upper and lower adjacent values and the line is the median. A Wilcoxon test was conducted for comparing the experiments and *** = p<0.001. Sample size: Control=234, Auxin=225. H) Scheme of the experiment in (I and J). I) The graph represents the percentage of EdU positive cells for HCT116:Ki-67-AID wt without (-) and with (+) Auxin and HCT116:Ki-67-AID STING KO without (-) and with (+) Auxin. The values are the mean of 2 biological replicas and the error bars represent the standard deviations. Samples size: HCT116:Ki-67-AID (-)Auxin=224, (+)Auxin=243, HCT116:Ki-67-AID STING KO (-)Auxin =290, (+)Auxin=236. The data were statistically analysed with a Chi-squared test. **= p< 0.01, ***= p<0.001. J) Distribution of the replicating cells according to the patterns shown in (Figure 1 G) from the experiment in H-I. The data were statistically analysed with a Chi-squared test. ***= p<0.001, ns= not significant.

One aspect that has emerged by studying Ki-67, is that different outcomes on Ki-67 silencing have been reported in different cell lines [27]. We therefore wanted to evaluate if the interferon pathway activation was a common theme, or it was dependant on p21 or p53 status. As there are no other cell lines available that have the endogenously AID tagged Ki-67, we used RNAi with two published Si-oligos [8] [11]. RNAi experiments were conducted in the HCT116*^Ki-67-AID^* (cell line we have generated), the original HCT116 carrying Ki-67 wt alleles, an HCT116 cell line where the MHL1 *wt* gene has been re-introduced (to note that HCT116 have a mutation in the DNA repair enzyme MHL1), SW840, a p53 negative colorectal cancer (HCT116 are p53 *wt*), HeLa and an immortalised non-cancerous cell line hTERT-RPE1. In all the cell lines used, and with both oligos we could reproduce the activation of the interferon response upon Ki-67 RNAi (Figure 3 D, E and Supplementary Figure 2 E-I). This clearly indicates that, without Ki-67, the cytosolic DNA-sensing immune response pathway is activated. However, as discussed before, we could not detect increased γH2AX in these cells (Figure 2 A), therefore we set out to check the presence of damage by other methods. We used the comet assay that assesses the presence of DNA damage directly. Cells where Ki-67 was degraded at the G1/S boundary showed a significant increase in DNA damage as visualised by the increased length of the comet tail (Figure 3 F, G).

We then asked the question if the interferon pathway activation could play a role in the S phase progression delay we observed upon Ki-67 degradation. To address this, we knocked out STING in the Ki-67-AID cell line, following the strategy described by Langeris et al. [31] (Supplementary Figure 5 A, B). Using this cell line, we blocked the cells in thymidine, degraded Ki-67 for 4 h and then released them from the block. We analysed both the ability of the cells to enter S phase and S phase progression in terms of EdU pattern distributions. The results obtained indicated that indeed the interferon pathway activation contributes to the delay of S phase progression with a pronounced effect on the replication timing for cells that enter S phase (Figure 3 H, I).

Overall, these data suggest that Ki-67 is important at the G1/S transition to protect the replication machinery. Lack of Ki-67 leads to the collapse of the replication machinery, persistence of DNA damage that is sensed by the cGAS-STING pathway and activates interferon responsive genes. This pathway also contributes to the observed delay in replication.

Interestingly, the expression of IFIT1 (interferon induced gene), STAT2 (Signal transducer and activator of transcription that mediates signalling by type I interferons) and Ki-67 are also negatively correlated in several cancer types as indicated by the analyses we have conducted using the R2 genomic platform (Supplementary Figure 5 C, D), suggesting that this pathway could be of more general relevance and should be taken into consideration when stratifying cancer types based on Ki-67 expression.

### Ki-67 interacts with components of the replication machinery

To gain a molecular understanding on the role of Ki-67 during DNA replication, we set out to conduct proteomic experiments. We wanted to capture Ki-67 interactome at the G1/S boundary specifically, and not relying on antibodies of solubilisation problems. To achieve this goal, we generated another cell line in HCT116 where we endogenously tagged Ki-67 with mClover and APEX2 (Figure 4 A and supplementary Figure 3 A, B) [32]. In this cell line, addition of biotin to the culture and a brief treatment with H_2_O_2_ leads to the biotinylation of proteins in close contact of Ki-67. The biotinylation can be visualised by TXRed-streptavidine where the nuclear space, some nucleoli regions and the chromosome periphery become highlighted (Figure 4 B). Using this cell line, combined with Stable Isotope Labelling with Amino acids in Cell culture (SILAC) approach, we labelled the Ki-67 interactome at the G1/S transition with biotin, purified the proteins with streptavidin column and subjected to mass spectrometry (Figure 4 C). STRING analyses of the protein list interactome showed a significant increase for proteins present in nuclear speckles, proteins involved in transcription, immune system processes and DNA conformational changes (Figure 4 D). Further Reactome-GSA analyses also highlighted the presence of proteins involved in DNA replication and DNA repair (Figure 4 E). Among them, the DNA helicase MCM3 was highly enriched, so we tested the possible interaction both by blotting the APEX pull down and by proximity ligation assay (PLA). We could detect MCM3 in the APEX pull down from cells at the G1/S boundary but not from cycling cells or control cells (Figure 4 F). Similarly, in the PLA assay, we could detect PLA signals between MCM3 and Ki-67 but not upon Ki-67 degradation (Figure 4 G, H). These experiments strongly support Ki-67 being in proximity of the replisome at the G1/S boundary.

**Figure 4.**
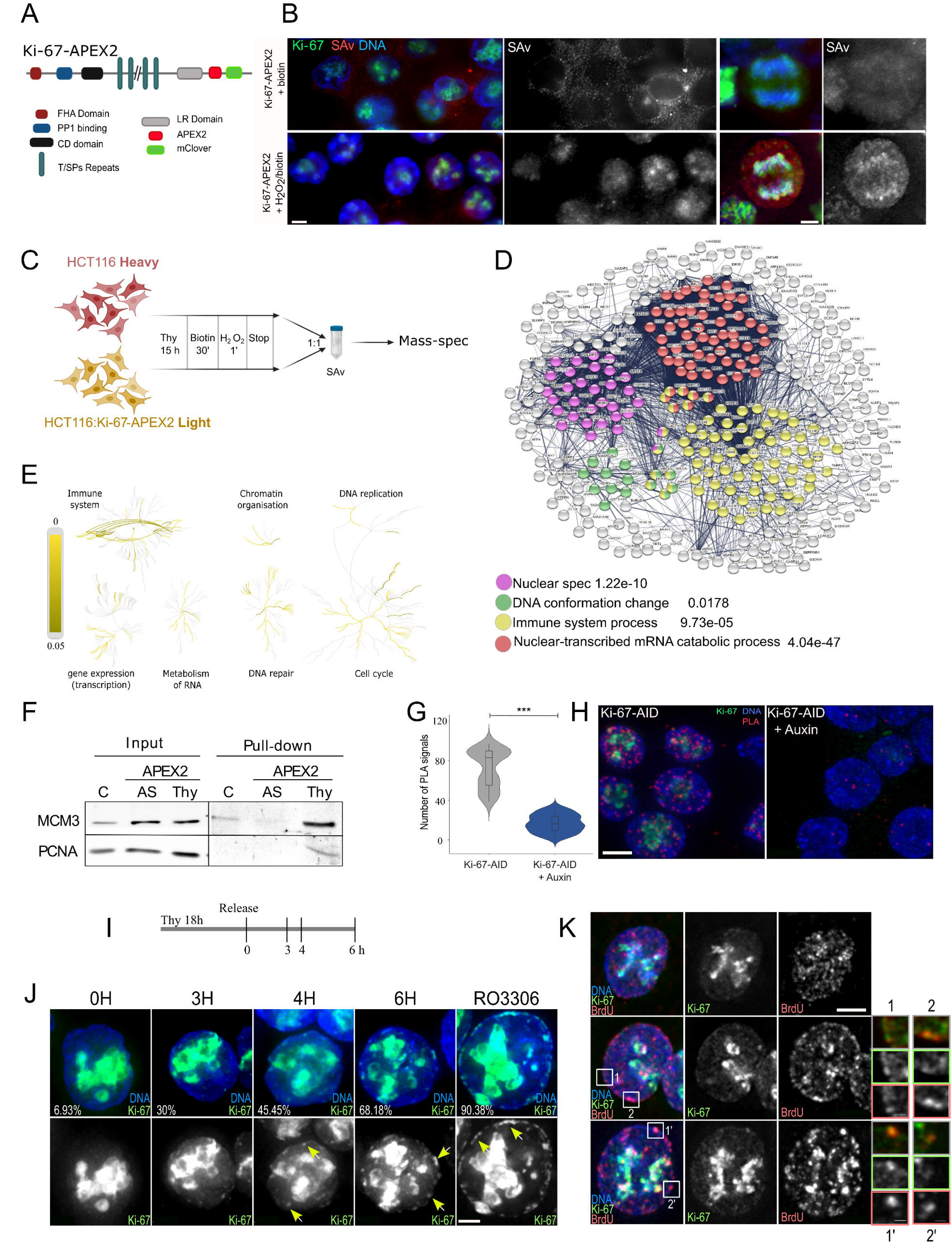
Ki-67 is in proximity of the replication machinery during S phase. A) Scheme of Ki-67-APEX2 tagged protein. B) Representative images of the HCT116:Ki-67-APEX2 cell line treated with biotin for 30 minutes and then with or without H_2_O_2_ for 1 minute. The cells were fixed and stained with RFP-streptavidin and DAPI. Scale bars 5 μm. C) Scheme of the SILAC experiment. Drawing made with Biorender. D) STRING analyses of the APEX proteome from the experiment in (C). E) Reactome pathway enrichment analyses of the Ki-67 interacting proteins form the experiment in (C). F) Western blot of HCT116 (C) or HCT116: Ki-67-APEX2 (APEX2) probed with antibodies against MCM3 (top panels) and PCNA (bottom panels). G) Violin plot of the quantification of the experiment in (H). The box inside the violin represents the 75^th^ and 25^th^ percentile, whiskers are the upper and lower adjacent values and the line is the median. Sample size: Control=114, Auxin=99. A Wilcoxon test was conducted for comparing the experiments and *** = p<0.001. H) Representative images of the proximity ligation assay (PLA) using anti MCM3 and anti GFP antibodies on HCT116:Ki-67-AID cell line without (left) of with (right) Auxin. Scale bar 5 μm. I) Scheme of the experiment for (J) and (K). J) Representative images of HCT116:Ki-67-AID cell line at different time points from the thymidine block release. The numbers represent the percentage of cells showing Ki-67 appearing as foci at the nuclear periphery. Cells were stained with DAPI. The yellow arrows indicate the foci. Scale bar 5 μm. K) Representative images of HCT116:Ki-67-AID cell line at early (top panels), middle (middle panels) and late (bottom panels) replication stages and stained with anti BrdU antibodies and DAPI. Scale bar 5 μm. 1 and 2 boxed regions in the left panels represent the enlarged images on the right. The frame colour of the enlargements indicates the specific channel.

We were very intrigued by these findings, and we decided to look more closely at the localisation of Ki-67 in replicating cells taking advantage of the cell line with the endogenously tagged protein. Upon release from the thymidine block, Ki-67 starts appearing in small foci at the nuclear periphery that grow in number and intensity throughout replication and reach a maximum of nuclear rim localisation in cells at the G2/M transition (Figure 4 I, J). Interestingly, heterochromatin and the nuclear periphery replicates middle-late, therefore we wondered if those appearing Ki-67 foci were the regions where DNA replication was occurring. Indeed, several of these foci were also positive for BrdU when added for 1 h before fixation (Figure 4 K and Supplementary Figure 3 C, D).

In summary, we conclude that Ki-67 interacts with components of the replication machinery at the G1/S boundary and its degradation leads to destabilisation of the replisome causing DNA damage that triggers interferon signalling, but it is not detected by the canonical DNA damage response.

### Ki-67 is important for the DNA damage response signalling by regulating Huwe1 activity

We next wondered why DNA damage could not trigger the expected signalling cascade when Ki-67 was degraded. We therefore re-analysed the Ki-67 interactome with a particular focus on proteins that interact with MCM3 and are enriched in Ki-67 APEX2 pulldowns. One of these proteins is the ubiquitin ligase HUWE1 which has also been shown to be present at the replication forks [33, 34] (Figure 5 A). Interestingly, it has been observed that the block of HUWE1 activity mediated by ATM and SIRT6/SNF2H is necessary for γH2AX foci formations at double strand breaks [35]. HUWE1 is a multi-faceted E3 ubiquitin ligase of the HECT family with many confirmed substrates, but mechanistic understanding of its functional roles in signalling remains limited. Amongst its known substates that could be relevant for this cell cycle stage are H2AX [36] and TP53 [37] (Figure 5 B).

**Figure 5.**
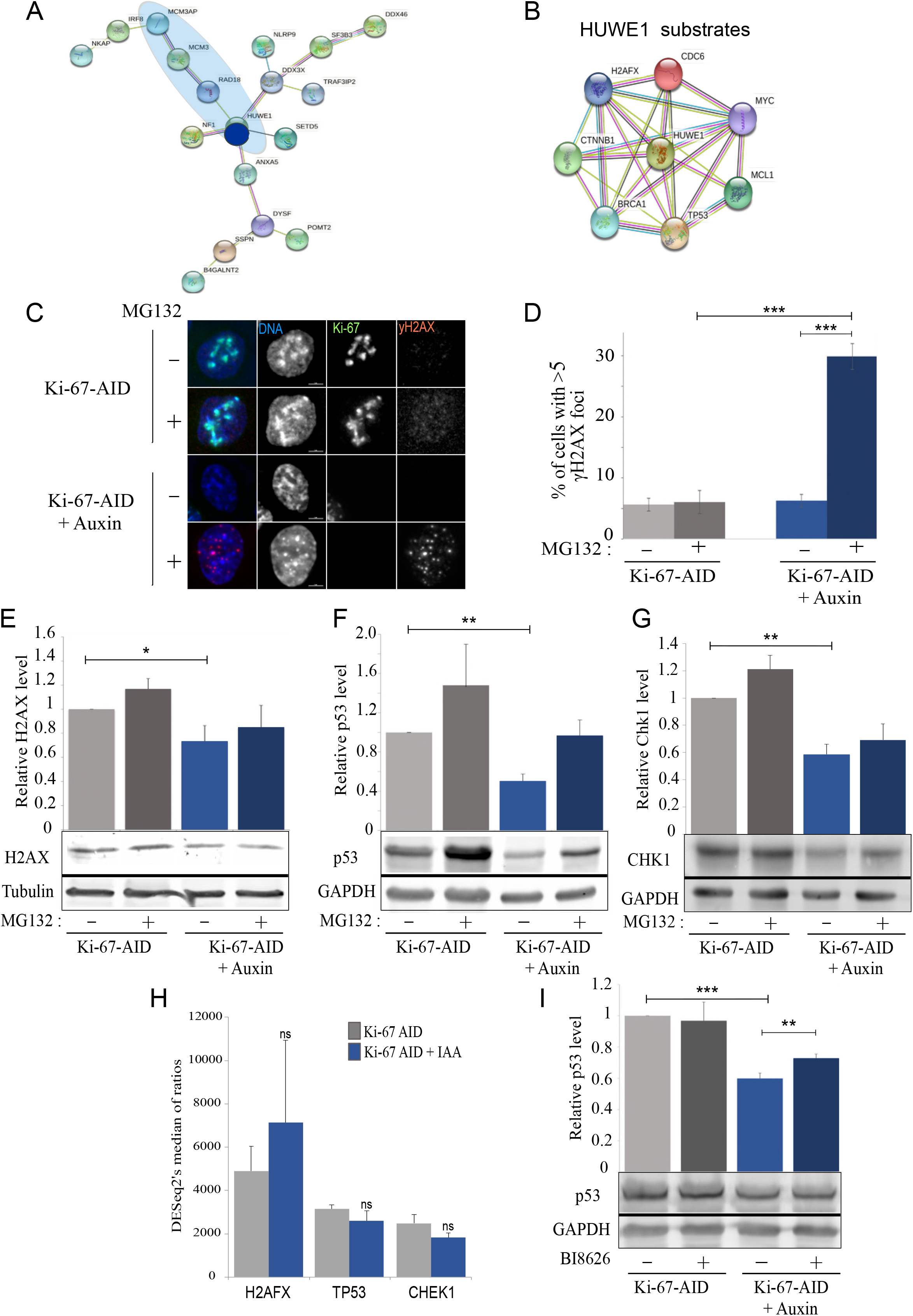
Ki-67 degradation leads to Huwe1 activation. A) STRING analyses of proteins form the APEX2 proteome that are linked to DNA replication (circled in blue). B) STRING representation of known HUWE1 interactors. C) Representative images of HCT116:Ki-67-AID cell line blocked with thymidine for 18 h then untreated (Ki-67-AID) or treated with auxin for 4 h (Ki-67-AID Auxin) in the presence (+) or absence (-) of MG132. The cells were fixed and stained for γH2AX (red) and counterstained with DAPI (blue). Sample size: Control=335, Control (MG132)=332, Auxin=334, Auxin (MG132)=311. Scale bar 5 μm. D) Quantification of the experiment in (C). The values represent the average of 3 independent replicas and the error bars are the standard deviations. The experiments were analysed by Chi-squared test. ***= p< 0.001. E-G) Representative Western blot analyses of HCT116:Ki-67-AID cell line blocked with thymidine for 18 h then untreated (Ki-67-AID) or treated with Auxin for 4 h (Ki-67-AID Auxin) in the presence (+) or absence (-) of MG132. The blots were probed with anti alpha tubulin or anti GAPDH antibodies and with anti H2AX (E), p53 (F) and CHK1 (G). The images were acquired with a LICOR machine in the linear range for quantification purposes. The graphs at the top represent the quantification of the blots. The values represent the average of 3 independent replicas and the error bars are the standard deviations. The experiments were analysed by a Student’s t-test. *=p<0.05 **=p<0.01. H) Expression levels of the indicated genes obtained from the RNA seq experiments described in Figure 3 A. The values represent the average of the 3 independent replicas and the error bars are the standard deviations. The experiments were analysed by a Student’s t-test. ns=not significant. I) Representative Western blot analyses of HCT116:Ki-67-AID cell line blocked with thymidine for 18 h then untreated (Ki-67-AID) or treated with Auxin for 4 h (Ki-67 AID Auxin) in the presence (+) or absence (-) of BI8626. The blots were probed with anti GAPDH and anti p53 antibodies. The images were acquired with a LICOR machine in the linear range for quantification purposes. The graph at the top represents the quantification of the blots. The values represent the average of 3 independent replicas and the error bars are the standard deviations. The experiments were analysed by a Student’s t-test test. **=p<0.01 ***= p< 0.001.

We hypothesised that the lack of γH2AX foci upon Ki-67 degradation could reflect the fact that H2AX was not stabilised. We therefore quantified the number of γH2AX foci in cells arrested in thymidine with and without Ki-67 and in the presence of the proteosome inhibitor MG132. These analyses revealed that cells without Ki-67 do not show an increase in γH2AX foci (even if there is more DNA damage (Figure 3 F, G)) but, upon treatment with MG132, the number of γH2AX foci increases significantly (Figure 5 C, D) (to be noted that the short MG132 treatment does not restore Ki-67 levels in cells – Figure 5 C). To test if this was due to H2AX degradation, we analysed the total level of this histone by western blot in thymidine arrested cells with and without Ki-67. The data show that indeed H2AX levels are reduced without Ki-67, and they are rescued by blocking the proteosome (Figure 5 E); this is not caused by a decrease of its mRNA (Figure 5 I). A similar behaviour was observed for other known HUWE1 substrates (Figure 5 F-G). Finally, we confirmed that HUWE1 seems to be the hyperactive ligase triggered by Ki-67 degradation because, by blocking its activity with the specific inhibitor BI8626, we observed increased TP53 and CHK1 levels (Figure 5 I and Supplementary Figure 4 A,B).

Altogether, these data indicate that lack of Ki-67 during S phase leads to the silencing of the canonical DNA damage response in a HUWE1/proteasome dependant manner.

### Ki-67 degradation impairs the phospho-regulation of several pathways

Cells lacking Ki-67 have a delay in S phase progression but do not enter apoptosis. This is quite an unusual signalling and we wanted to understand the molecular cascade that is at the basis of this phenotype in an unbiased manner.

We used again a SILAC based proteomic approach; we blocked the cells with thymidine and then degraded Ki-67 in “light” labelled samples. This time we isolated the nuclei and processed the samples for phospho-proteomic to identify changes in the signalling pathways (Figure 6 A).

**Figure 6.**
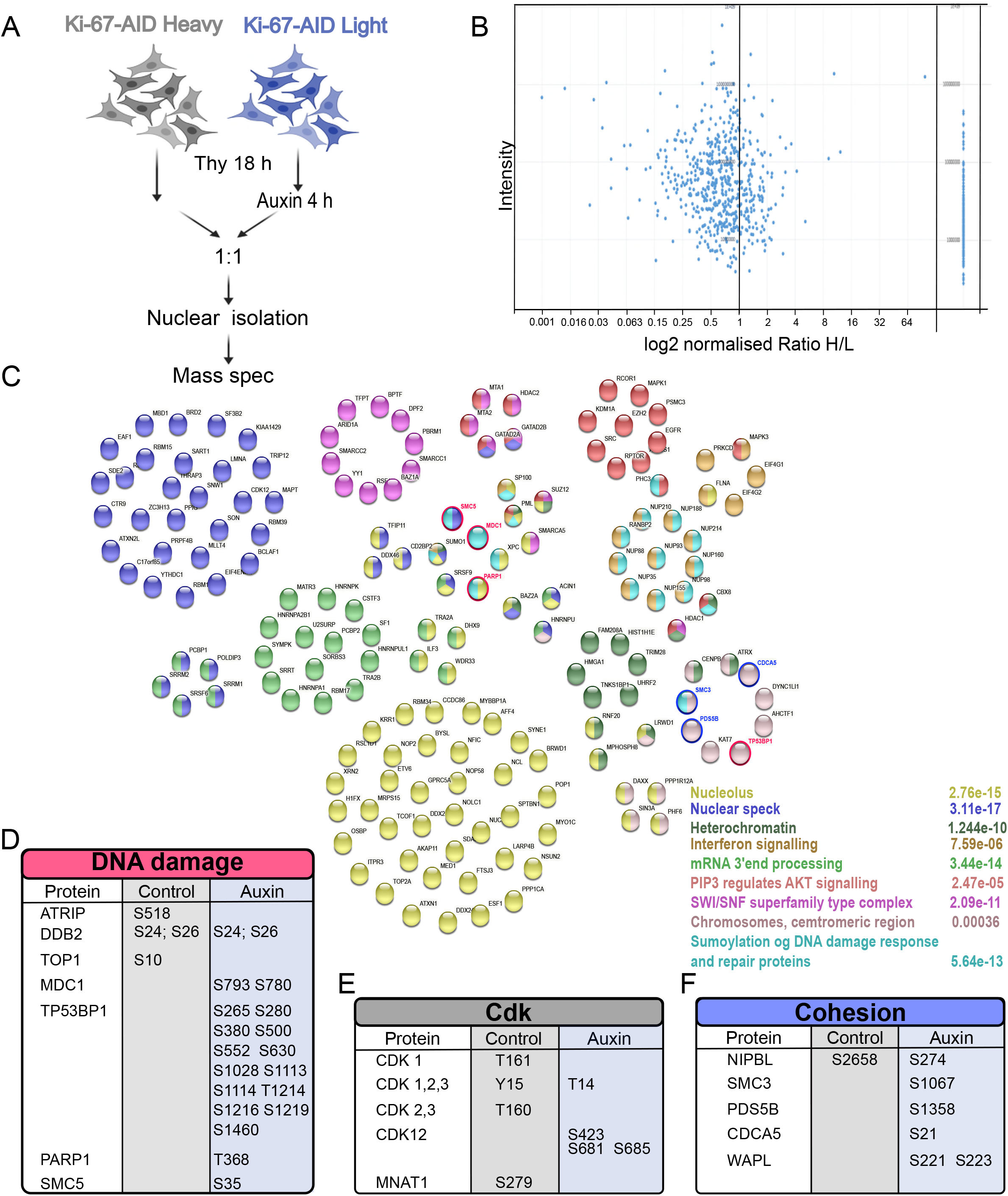
Changes in the phosphoproteome upon Ki-67 degradation at the G1/S boundary. A) Scheme of the experiment. B) Graphs representing the log2 ratio between heavy and light peptides and their normalised intensity. On the right are represented the proteins that were only found phosphorylated in the light sample (HCT116:Ki-67-AID + Auxin). C) Representations of the different protein groups enriched in the HCT116:Ki-67-AID + Auxin phosphoproteome. The different colours relate to the named categories and the numbers represent the false discovery rate. The red and the blue circles highlight the DNA damage-related proteins and Cohesin/regulator proteins respectively. D-F) List of the proteins and their phosphosites enriched in the HCT116:Ki-67-AID (Control) and HCT116:Ki-67-AID + Auxin (Auxin) samples.

The analyses revealed that several proteins were hyperphosphorylated upon Ki-67 depletion while only a smaller number were underphosphorylated compared to the control (Figure 6 B). STRING analyses of the hyper-phosphorylated proteins highlighted different categories (Figure 6 C). A good proportion of the proteins were nucleolar components: this should not come to a surprise since Ki-67 itself is a nucleolar protein. The second major group was comprised by nuclear speckle proteins; this group had already been identified in the APEX2 proteomic (Figure 4 D) and again demonstrates a close link between Ki-67 and these nuclear structures. A similar result was obtained for a group of proteins involved in 3’mRNA splicing. The interferon signalling pathway was enriched as well, and this further strengthens the link between lack of Ki-67 and the innate immune response, possibly triggered by DNA damage. Indeed, some DNA damage related proteins were present in the hyperphosphorylated proteome (Figure 6 D). Phosphorylation of MDC1 and TP53BP1 were only found upon Ki-67 degradation. These phosphorylations indicate activation of MDC1 and TP53BP1 in response to DNA damage [38] [39]. Interestingly, this pathway seems to function either in regions where H2AX is not present or in conditions where the γH2AX pathway is not working; our data of undetectable γH2AX foci upon Ki-67 degradation at the G1/S boundary (Figure 2 A and Figure 5 C, D) are in agreement with the latter [40].

These analyses also revealed that CDKs regulation is quite different in the two conditions: CDK2,3 is phosphorylated at T160, which represents an activating phosphorylation [41] but, at the same time, CDK1,2,3 is also phosphorylated at Y15, which is an inhibitory phosphorylation [41]. CDK2 Thr160 phosphorylation increases during S and G2 when CDK2 is active. Tyr15 phosphorylation has been observed upon replication stress (stalled replication forks) and has been linked to the activation of the WEE1 AND MYT1 kinases as DNA damage response signalling [42]. Upon Ki-67 degradation, CDK 1,2,3 is (are) phosphorylated at Thr14, which is another inhibitory phosphorylation. Altogether, these results strongly suggest that, in Ki-67 depleted cells, CDK2 is inactivated (Figure 6 E). We tested this hypothesis using a reporter for CDK2 activity [43] [44]. The sensor includes amino acids 994–1087 of human DNA helicase B fused to mCherry, and contains four CDK consensus phosphorylation sites, a nuclear localization signal, and a nuclear export signal. In S phase it has an intermediate activity (localises both to the nucleus and the cytoplasm) while in G0/G1 is inactive (localises only to the nucleus). We generated a HCT116*^Ki-67-AID^*stable cell line expressing the mCherry-CDK2 reporter. Cells were blocked in Thymidine as described before, and the localisation of the reporter was analysed in cells with and without auxin treatment. The ratio between the nuclear and cytoplasmic localisation of the reporter shows a significant decrease in CDK2 activity upon auxin treatment that explains the block or delay in resuming replication after release (Supplementary figure 4 C, D). Therefore, we conclude that CDK2 is inactivated upon Ki-67 degradation.

Other chromatin remodelling and structural proteins were also found to be hyperphosphorylated in auxin. These belong mainly to 3 categories: the SWI/SNF complex, heterochromatin and centromeric chromatin. The SWI/SNF complex has been linked to DNA repair mechanisms as well as transcription regulation and R loop resolution [45].

A closer inspection of the hyperphosphorylated proteins that belong to the cluster of “chromosome, centromeric region” revealed that many components are linked to cohesion biology namely SMC3, WAPL, CDCA5, NIPBL (Figure 6 F). This is quite interesting, in fact, during replication, cohesion establishment is very important for genome maintenance [46]. Interestingly, some of the phosphorylation sites in SMC3 have been shown to increase the binding to WAPL that functions in removing cohesion form chromatin. This process could be linked to WAPL-dependent repair of damaged DNA replication pathway that has recently been proposed, whereby increased cohesin removal is required to complete DNA synthesis under conditions of persistent DNA replication stress [47].

In conclusion, these analyses provide us with a picture whereby Ki-67 degradation leads to inactivation of CDK2, triggers a DNA repair pathway independent of γH2AX, activates an interferon response and alters cohesion regulation.

### Ki-67 degradation causes replication and sister chromatids cohesion defects

As shown by the growth curves (Figure 1 D), cells without Ki-67 can eventually resume the cell cycle, although the growth rate is much slower. Therefore, we investigated if, given the time, they can override the replication block. For this, we measured the percentage of G2 cells that have replicated the DNA after release from thymidine block in BrdU and RO3306 (a CDK1 inhibitor) (Figure 7 A). This treatment would capture cells that completed S phase and arrested at the G2/M boundary. The analyses indicated that most (83%) of the cells where Ki-67 had been degraded could complete replication during this time, a proportion that is not significantly different from the controls (Figure 7 B-C). This provided us the opportunity of testing how replication occurred at different genomic regions and the status of sister chromatid cohesion.

**Figure 7.**
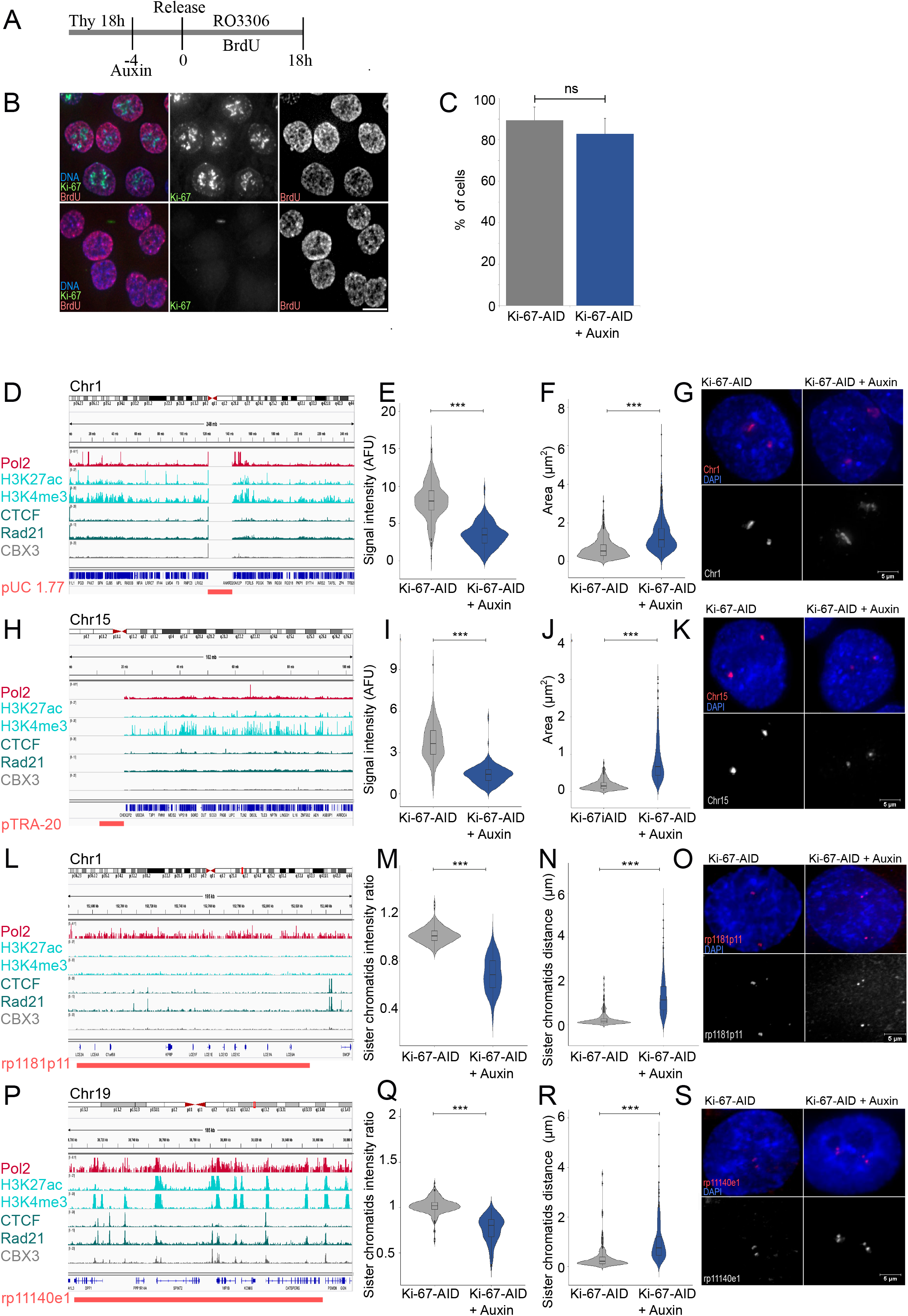
Ki-67 degradation in S phase causes replication and sister chromatid cohesion defects. A) Scheme of the experiment. B) Representative images of the experiment in (A). Cells were fixed at the end o the experiment and stained with anti-BrdU antibodies. Sample size: Control=326, Auxin=345. Scale bar 5 μm. C) Quantification of the experiment in (B). The data were analysed with a Chi-squared test. ns= not significant. D,H,L,P) IGV tracks of the regions selected for FISH analyses showing the chromosome position, Pol2, H3K27ac, H3K4me3, CTCF, Rad21 and CBX3 ChIP seq profiles form Encode and obtained from experiments in HCT116 cells. Magenta lines at the bottom represent the location of the probe selected for the satellite of chromosome 1 (D), satellite of chromosome 15 (H), single copy low expression region on chromosome 1 (L) and single copy high expression region on chromosome 19 (P). E, I) Quantification of the intensity of the FISH signals in the HCT116:Ki-67-AID and HCT116:Ki-67-AID + Auxin samples for the probes in D (E) and H (I). Sample size: (E): Control=279, Auxin= 285, and (I): Control=309, Auxin=274. F, J) Quantification of the area occupied by the FISH signals in the HCT116: Ki-67 AID and HCT116: Ki-67 AID + Auxin samples for the probes in D (F) and H (J). M, Q) Ratio of the FISH signals intensity between the two sister chromatids in the HCT116: Ki-67 AID and HCT116: Ki-67 AID + Auxin samples for the probes in L (M) and P (Q). Sample size: (M): Control=214, Auxin=205, and (Q): Control=214, Auxin=237. N, R) Distance of FISH signals between the two sister chromatids in the HCT116: Ki-67 AID and HCT116: Ki-67 AID + Auxin samples for the probes in L (N) and P (R). Sample size (N): Control=2, Auxin=246 and (R): Control=207 Auxin=220. Al the data presented in the violin plots were statistically analysed with a Wilcoxon test. ***=p<0.001. The box inside the violin represents the 75^th^ and 25^th^ percentile, whiskers are the upper and lower adjacent values and the line is the median. G, K, O, S) Representative images of the Fish signals obtained in HCT116:Ki-67-AID and HCT116:Ki-67 -AID + Auxin samples for the probes in D (G), H (K), L (O) and P (S). FISH signals in magenta and DNA in blue. Scale bar 5 μm.

We selected 4 different genomic regions: 1) the satellite of chromosome 1 as it represents a large heterochromatic region (Figure 7 D); 2) the satellite of chromosome 15 as it is a nucleolar organiser region (NOR) carrying chromosomes (that contributes to the nucleolar function) (Figure 7 H); 3) a region that is not transcribed on chromosome 1 (as shown in the RNAseq datasets we conducted and supported by the ChIPseq data in UCSC) (rp1181p11) (Figure 7 L) and 4) a region that is highly transcribed on chromosome 19 (rp11140e1) (Figure 7 P).

We then conducted FISH experiments on nuclei of cells that went through replication with and without Ki-67 and were then arrested at the G2/M boundary with RO3306.

For the satellite probes, we measured the total intensity of the signal in the nuclei and the area occupied by each signal. These analyses revealed that the probe signals in cells without Ki-67 was significantly diminished (Figure 7 E, G, I, K) but the area occupied was larger (Figure 7 F, G, J, K) indicating that replication did not occur completely, and that chromatin organisation was also impaired.

For the single copy probes, we analysed the intensity of the signal of each sister chromatid in the pair: if replication occurs correctly the ratio of intensity between the two sisters should be 1; if replication does not occur precisely, you could expect some deviations from the 1:1 ratio depending on where the replication fails. We also measured the distance between the sister chromatids. As the data show, Ki-67 degradation causes a change in the 1:1 intensity ratio (Figure 7 M, O, Q, S) but also leads to an increase in sister chromatids distance for both loci (Figure 7 N, O, R, S), although the latter seems more prominent for the non-transcribed locus than for the transcribed one.

These experiments reveal for the first time that depletion of Ki-67 at the beginning of S phase causes defects in DNA replication and impairs sister chromatid cohesion. In this respect, it would be interesting to evaluate how these problems contribute to the mitotic phenotype that has been reported for Ki-67 depletion and how cell adapt to a prolonged absence of Ki-67.

## DISCUSSION

In this study we have addressed the role of KI-67 in cell cycle progression with a particular focus at the transition from G1 to S. Despite Ki-67 being the election marker for cell proliferation, several studies seem to suggest that it is dispensable for it [11] [13]. However, cells without Ki-67 appear to have a phenotype, including reduced ability of metastasise in xerographs implants [15] and, in the case of the Ki-67 knock-out mice, being resistant to chemical or genetic induction of intestinal tumorigenesis [19]. For these reasons and the link to its prognostic value in several cancers, it is important to conduct a careful analysis of how cell cycle transitions are affected by Ki-67. Detailed analyses of Ki-67 function at a molecular level have mainly focussed on mitosis. Important studies on phenotypes related to different cancer types have not reached yet the level of mechanistic insights necessary to understand if this protein could also lead to therapeutic avenues.

We have therefore generated new HCT116 cell lines to achieve both a homogeneous and rapid degradation of Ki-67 in a short time period (2-4 h) using the AID-auxin degron strategy [48] and to identify interacting proteins using the APEX2 technology [32]. When cells are maintained in presence of auxin, growth and progression through S phase are delayed. This observation per se is already important and reconcile some contrasting experimental evidence in the literature about the need of Ki-67 for cell proliferation. In fact, a short treatment or RNAi could indeed show a cell cycle delay, but the generation of cell lines lacking Ki-67 would provide the window of opportunity for the cells to overcome the block and adapt to the new homeostasis.

Ki-67 degradation at the G1/S transition causes severe delays in resumption of DNA replication and progression through the replication stages. The delay occurs when either cells are arrested with thymidine or with the CDK4/6 inhibitor Palbociclib abut also when cells are already replicating and without the addition of any drug, thus confirming a role for Ki-67 during DNA replication. This observation agrees with data obtained in hTERT-RPE1 cells where depletion of Ki-67 by RNAi delayed S-phase entry upon arrest with hydroxyurea [27]. However, while the study by Sun et al [27] reported that the delay was caused by downregulation of genes related to DNA replication and dependent on p21 induction, our study provides a different mechanism. We have conducted RNA-sequencing experiments upon degradation of Ki-67 in cells arrested at the G1/S transition and we did not observe changes in transcription any replication factors. We have shown that transcriptional changes of replication components indeed occur when Ki-67 is degraded before mitotic exit but not at the G1/S transition; therefore, the changes in the transcriptional landscape observed using siRNA approaches are possibly a secondary effect of cells exiting mitosis without Ki-67 (Supplementary Figure 2 B). We also did not observe an increase in p21 levels, actually we have seen a decrease in p21 and we did not detect alterations in several chromatin markers, suggesting that epigenetics (at least for the markers we tested) is unlikely to be the source for the delay. Our analyses indicate a more direct role of Ki-67 as interactor of the replication machinery. The replisome appears to be unloaded upon Ki-67 degradation which can explain the delay in DNA replication; moreover, we could detect Ki-67 in close proximity of the MCM3 helicase. This is the first time that Ki-67 has been linked directly to the control of replication. Interestingly, and taking advantage of the endogenously tagged cell line, we could follow Ki-67 localisation during replication, and we could see changes in its localisation pattern as DNA replication progresses with Ki-67 co-localising with the replication foci, particularly evident for the late replicating regions of the genome: this aligns partially with a recent report showing that Ki-67 interacts with of Lamin B1-depleted late replicating genomic regions [28].

Despite observing a replication delay, a collapse of the replisome and inactivation of CDK2, we did not see the expected signalling pathway that could be triggered by DNA damage [49]: no p21 increase (the level were reduced compared to the controls) and no γH2AX foci. This picture was quite unusual and puzzling and led us to look in more detail at the APEX2 proteome. We identified the ubiquitin ligase HUWE1 as one of the interactors captured by biotinylation. Interestingly, HUWE1 has been shown to be present at the replication forks [33] [34] (Figure 5 A) and the block of HUWE1 activity mediated by ATM is necessary for γH2AX foci formations at double strand breaks [35]. This block does not seem to happen when Ki-67 is degraded during replication, leading to p21, p53 and H2AX degradation (all HUWE1 substrates [36, 37]), thus providing a molecular explanation for the lack of the canonical signalling pathway.

H2AX is highly expressed in actively growing cells, but it is usually downregulated in normal cells in quiescence. In fact, normal cells generally enter a growth-arrested state with marked downregulation of H2AX in vivo and in vitro [35] but also show increased H2AX proteasomal degradation mediated by HUWE1 [35]. Ki-67 is also not expressed in quiescent cells and the parallelism with our new findings during replication is quite intriguing; it would be interesting in the future to understand more about the involvement of Ki-67 in entering, maintaining, and exiting the quiescent state.

Although cells lacking Ki-67 delay their progression to S phase, they eventually manage to survive: how is this possible? We therefore looked at the changes in signalling that occur at this stage of the cell cycle upon Ki-67 degradation by conducting a SILAC based phospho-proteomic. This approach led us really to have a detailed picture of the signalling cascade. When Ki-67 is degraded, there is hyperphosphorylation of MDC1 and TP53BP1; this pathway seems to be important for DNA repair either in regions where H2AX is not present or in conditions where this histone cannot be phosphorylated [40]. This finding further reinforces that lack of Ki-67 triggers an alternative DNA damage repair pathway. Moreover, upon Ki-67 degradation, we also clearly see CDK2 inactivation. These data further demonstrate that Ki-67 degradation causes the collapse of the replication forks and DNA replication stress.

Other important phosphorylations only present after Ki-67 degradation were found on components of the cohesin complex and cohesion loading and unloading components. These suggest that cohesin is possibly removed from the chromatin to facilitate the repair process. In fact, in untransformed cells, replication stress triggers sister chromatid cohesion loss mediated by the cohesin remover WAPL [47]. Consequently, cells that transit through replication after Ki-67 is degraded show cohesion defects in G2.

We therefore propose that Ki-67 has an important role during replication and helps to coordinate timely replication progression and forks protection thus proving to be an important player in genome stability. These data could also explain the link between Ki-67 levels and sensitivity to CDK inhibitors treatment [24].

Ki-67 has been shown to bind PP1 [8] but not many substrates have been identified apart from Ki-67 itself [50]. We therefore wondered if the phenotype observed by Ki-67 degradation was mediated by PP1. If this was the case, we would expect to find some of hyperphosphorylated proteins (real substrates) also present in the APEX proteome. However, the overlap of the two datasets only presented few common components (ANXA2P2, ANXA2, ARHGEF2, B4GALT1, DDX46, DSP, HNRNPH1, HNRNPH2, MVP, POLDIP3, PPIG, RBM39, U2SURP), which are unlikely to represent the key triggers for all these events; however, it will be interesting to evaluate if they represent real and novel Ki-67/PP1 substrates. We therefore tend to suggest that the effect of Ki-67 in DNA replication is likely to be PP1 independent.

Although we did not observe major changes in transcription after Ki-67 degradation at the G1/S transition, we noticed that some genes related to the interferon response were upregulated. This could be a consequence of the collapsed replication forks where DNA fragments are recognised by STING, thus leading to the interferon response. The activation was also confirmed by phosphorylation of proteins involved in this signalling cascade. The interferon pathway activation is not restricted to the cell line we used, and the same response can be triggered via RNAi approaches and in different cell lines. This pathway seems also to contribute to the delay in S phase progression observed upon Ki-67 degradation since the knock-out of STING significantly improves the progression of replication even in the absence of Ki-67.

Because Ki-67 expression is an important prognostic marker in several cancer types, we have interrogated the R2 genomic platform and correlated Ki-67 expression to IFIT1 and STAT2 (both belonging to the interferon pathway that are upregulated upon Ki-67 degradation). Interestingly, the two genes show a strong negative correlation among several dataset thus suggesting that the interferon response triggered by loss or low levels of Ki-67 protein is a common trend in several cancers and could be of importance in understanding tumour evolution and response to therapy.

Nevertheless, after the delay, cells where Ki-67 is chronically degraded or knocked out can still grow. Activation of the interferon pathway could explain some of the phenotypes observed in cancer cells where Ki-67 was knocked down such as reduced tumor growth in xenografts for HeLaS3 and abrogation of metastasis in the 4T1 model [19]. This aspect can be important in modulating the immune response towards cancer cells and could potentially represent a vulnerability pathway that can be explored. However, more studies will be required to really understand the long-term consequences and in vivo tumor evolution.

In summary, we have shown that Ki-67 is important for timely DNA replication and genome maintenance. The initial block leads to DNA damage that triggers an interferon response that could lead the cell to a new homeostasis but also can explain the less tumorigenic behaviour of Ki-67 knock-out cells. Interestingly, several studies [51] [52] [53] have demonstrated that mouse ESC-derived cells have limited response to inflammatory cytokines and various infectious agents and are deficient in type I IFN expression [54], therefore lacking an innate immune response. This scenario could also potentially explain why it was possible to obtain a mouse model lacking Ki-67 and bypass the initial delay stages of cell cycle and growth delay.

## Methods

### Experimental model and subject details

HCT116 cells were grown in Gibco™ McCoy’s 5A Medium GlutaMAX supplemented with 10% foetal bovine serum (FBS) and 1% Penicillin–Streptomycin (Gibco) at 37 °C with 5% CO2.

hTERT-RPE1 cells were grown in DMEM/F-12, GlutaMAX supplemented with 10% foetal bovine serum (FBS) and 1% Penicillin–Streptomycin (Gibco) at 37 °C with 5% CO2.

HeLa Kyoto and SW480 cells were grown in DMEM GlutaMAX supplemented with 10% foetal bovine serum (FBS) and 1% Penicillin–Streptomycin (Gibco) at 37 °C with 5% CO2.

The HCT116 OsTR1 stable expressing cell line was kindly provided by Dr Masato T. Kanemaki (University of Tokyo, Japan).

The HCT116(wt), HCT116 (wt, MLH1 rescued), and SW480 cell lines was kindly provided by Dr Anabelle Lewis (Brunel University of London, UK).

The hTERT-RPE1 cell line was kindly provided by Dr Viji Draviam (Queen Marry University of London, UK).

### Method Details

#### Plasmids

pX330-U6-Chimeric_BB-CBh-hSpCas9 (no. 42230, Addgene) plasmids containing guide RNAs designed with CRISPOR (http://crispor.tefor.net), were generated according to [55].

To generate the plasmid donor containing the Ki-67 homology arm, genomic DNA from HCT116 (gift from Prof. Kanemaki, Japan) was purified by using EchoLUTION CellCulture DNA Kit (BioEcho), and 1175bp genomic DNA region around the termination codon of MKi67 was amplified using Phusion High-Fidelity DNA Polymerase

(Thermo Fisher Scientific, Massachusetts, United States) (primers are listed in Supplementary Methods Table 1), cloned in pGEM-T Easy empty vector (Promega) and sequenced. Based on sequencing, the plasmid containing MKi67 homology arms was synthetized, the stop codon was mutated into a BamHI restriction site, and point mutations in PAM sequences. The mAID-mClover-HygR cassette or mAID-mClover-NeoR cassette [48] was inserted into the BamH1 site in with the MKI67 to generate the homology arms donor plasmids. APEX2 was obtained by PCR from APEX2-csChBP (no. 108876, Addgene) and inserted in pMK-289 (no. 72827, Addgene) and pMK-290 (no. 72831, Addgene) using SacI/NheI restriction sites, thus replacing mAID by APEX2. pMK-289 and pMK-290 containing APEX2 were inserted between MKI67 homology arms at the BamHI site.

gRNAs are listed in the in Supplementary Methods Table 1.

#### Cell line generation

For the generation of Ki-67 AID and APEX2 cell lines, HCT116 cells were transfected with 2μg of total DNA of px330 cas9 and homology arms plasmids (mAID-mClover-HygR and mAID-mClover-NeoR for AID cell line and APEX2-mClover-HygR and APEX2-mClover-NeoR for APEX cell line) in 1:1:1 ratio and selected with 100 μg/mL hygromycin B (Invitrogen) and 700μg/ml of Geneticin (Gibco). Clones were first selected by microscopy and then screened via PCR genotyping using 3 different sets of Primers. The Primers are listed in the Table S1.

For the generation of HCT116:Ki-67-AID-DHB, HCT116:Ki-67-AID cells were co-transfected with 2μg of total DNA of CSII-pEF-hDHB-mCherry and pMSCV-Blasticidin in 9:1 ratio and selected with Blasticidin 10μg/ml (Sigma). Clones were selected by microscopy.

#### Transfections

For siRNA treatments, HeLa, HCT116:Ki-67-AID, HCT116(wt), HCT116 (wt, MLH1 rescued), and SW480 cells were seeded into 24-well plates, transfected using Polyplus JetPrime® (PEQLAB) with the appropriate siRNA oligonucleotides (50 nM) and analysed after 72 h. The siRNAs were obtained from Merck. The oligos are listed in the Table S1

#### Immunofluorescence microscopy

Cells were fixed in 4% PFA and processed as previously described [56]. Primary and secondary antibodies were used as listed in the key resources table. Three-dimensional data sets were acquired using a wide-field microscope (Delta Vision) Cascade II:512 camera system (Photometrics) and Olympus UPlanSApo 100x/1.40na Oil Objective (Olympus) and a wide-field microscope (NIKON Ti-E super research Live Cell imaging system) with a 100X Plan Apochromat lens, numerical aperture (NA) 1.45.

The data sets were deconvolved with the delta vision software or NIS Elements AR analysis software (NIKON). Three-dimensional data sets were converted to maximum projection, exported as TIFF files, and imported into Inkscape for final presentation.

#### Immunoblotting

Whole cell extracts were prepared by direct lysis in 1× Laemmli sample buffer [57]. Cellular fractionation was performed according to paper Herrmann, et al (Herrmann, Avgousti and Weitzman, 2017), separated in SDS–PAGE and transferred onto nitrocellulose membranes.

Membranes were blotted with primary and secondary antibodies, that are listed in the in Supplementary Methods Table 2. Membranes were visualised using either the Bio-Rad ChemiDoc XRS system or the LiCor Odyssey system.

#### Flow cytometry cell cycle analysis

Cells were trypsinised, resuspended and incubated at room temperature for 30 min in 70% ice-cold ethanol. Cells were centrifuged at 1,000 g for 5 min, washed with PBS and the supernatant discarded. The pellet was resuspended in 200μl of RNase A/PBS (100 μg/ml) and incubated for 2 h at 37°C in the dark. Propidium iodide (Fisher Scientific, P3566) was added at a final concentration of 5μg/ml just before analysing the samples by flow cytometry using the ACEA Novocyte Flow Cytometer. The analysis was performed using the NovoExpress® software.

#### S phase progression analyses

For S phase progression analyses with EdU incorporation, cells were treated with Doxycycline (2μg/ml) and 2 mM thymidine, 24 and 18 h before the addition of Auxin 1000μM, respectively. After the 4 h treatment with Auxin the cells were released and Edu added 30min before fixation at the and click it reaction was performed with Click-iT™ EdU Cell Proliferation Kit for Imaging, Alexa Fluor™ 647 (Thermo Fisher Scientific) according to the manufacturer protocol.

For BrdU incorporation analysis in G2, cells were treated with Doxycycline (2μg/ml) and 2 mM thymidine, 24 and 18 h before the addition of Auxin 1000μM, respectively. After the 4 h treatment with Auxin the cells were released in RO3306 and BrdU (Biolegent) added according with the manufacturer recommendation, and cells were fixed after 18 h.

For BrdU colocalization with Ki-67 foci during replication progression, cells were treated with 2 mM thymidine for 18 h. After 18 h the cells released and BrdU was added 1h before fixation according with the manufacturer.

For BrdU staining cells were permeabilised for 10min with 0.2% Triton-X, then 30min 2M HCL; the cells were then washed 3x 5min wash with PBS. 30min blocking at 37°C with 1% BSA was followed and finally 1h incubation with Alexa Fluor® 647 anti-BrdU. The cells were mounted after washing with PBS 3x 5 min.

#### CDK2 activity analyses

HCT116:Ki-67-AID-DHB cells were treated with Doxycycline (2μg/ml) and 2 mM thymidine, 24 and 18 h before the addition of Auxin 1000μM, respectively. After the 4 h treatment with Auxin the cells were fixed. Coverslips were mounted on DAPI and observed on the previously mentioned wide-field NIKON microscope. The analysis of CDK2 activity was conducted in ImageJ according to Cappell et al., 2016 (Cappell et al., 2016). Violin plots were generated with R Studio. The Wilcoxon statistical analysis was conducted with R Studio.

#### Comet assay

The alkaline comet assay was performed modified from [58]. Polylysine coated slides were first covered with high-melting-point 1% agarose and dried overnight at ambient temperature. After cell treatment, a drop of normal-melting-point agarose was first loaded on a slide, and then a drop of low-melting-point agarose was put on the precoated slide. Then 10^4^ cells were placed in the agarose droplet. Slides were lysed in a detergent solution (containing 2% N-Lauroylsarcosine sodium salt, 0.5M NA2EDTA and 0.1mg/ml proteinase K), for 1 hour at 4°C. DNA unwinding was carried out with an alkaline solution (90mM Tris Buffer, 90mM boric acid and 2mM NA2EDTA) for 90 minutes at room temperature in a 1-L electrophoresis unit. Then, electrophoresis was conducted for 40 minutes (20 V) in the same buffer solution. After the electrophoretic run, the slides were neutralized with distilled water, stained with SYBR Safe in NE buffer, dehydrated dipped respectively into 70%, 90% and 100% ethanol, with and dried overnight at room temperature. Images were captured using a Leica DM4000 fluorescence microscope (Leica Microsystems). Comet length was measured with ImageJ.

#### Luciferase reporter assay

24 h after siRNA knockdown transfection in 24-well plates as described for HeLa, HCT116:Ki-67-AID, HCT116(wt), HCT116 (wt, MLH1 rescued), and SW480, cells were co-transfected with plasmids expressing luciferase reporter constructs (IFNβ-firefly at 250ng and Renilla at 5 ng) (kindly provided by Prof Eric Schirmer, Edinburgh, UK). To measure luminescence produced by luciferase activity, cells were harvested 72h after the initial siRNA knockdown transfection. Luciferase assays were carried out with Dual-Luciferase Reporter Kit, Promega following manufacturer’s instructions. Light emission was measured with Junior LB 9509 (Berthold Technologies). Luminescence signal produced by IFNβ/NF-κB-firefly luciferase was divided by pRL-TK-renilla luciferase luminescence signal to control for variation in transfection efficiency and cell number.

#### Fluorescence in situ hybridisation (FISH)

HCT116:Ki-67-AID were treated with Doxycycline (2μg/ml) and 2 mM thymidine, 24 and 18 h before the addition of Auxin 1000μM, respectively. After the 4 h treatment with Auxin the cells were release in RO3306 for 18 h. The cells were fixed as follows: 15min 75mM Kcl, 3/1 MeOH/Acetic Acid ice cold for 30m (this step was repeated twice) finally the cells were stored in MeOH/acetic acid at −20°C. PAC-derived probes were extracted from bacterial cultures and fluorescently labelled by nick translation using Nick Translation Kit (Abbott) following the manufacturer’s instructions. The probe was separated using G-50 column, precipitated at −20°C in the presence of 10X salmon sperm DNA for the centromeric probes and salmon sperm and COT-1 DNA for single copy probes, according to the protocol of Garimberti and Tosi [59].

#### Pericentromeric/ a-satellite FISH

The probe (3μL of probe 7μL of hybridisation buffer - 50% (v/v) Formamide, 10% (w/v) dextran sulphate, 1x Denhart’s, 2× SSC, pH 7.0) was denatured at 73°C for 5. Probe and nuclei were denatured at 85°C for 5 min, and then hybridised overnight 39°C. Slides were washed with 2X SSC RT 10’, 0.4X SSC 60°C 10’, 0.1X SSC RT 10’ and counterstained with DAPI.

#### Single copy locus

The probe was added to the coverslips. Probe and nuclei were denatured at 75°C for 2min and then hybridised overnight at 37°C. The slides were washed in 2X SSC RT, 0.4X SSC at 72°C 5min, 2X SSC 0.05% Tween20 RT5 min and 1X PBS 5min and counterstained with DAPI.

For all FISH analyses, three-dimensional data sets were acquired using a wide-field microscope (NIKON Ti-E super research Live Cell imaging system) with 100X 1.45 (NA) Plan Apochromat lens. The data sets were deconvolved with NIS Elements AR analysis software (NIKON). Three-dimensional data sets were converted to Maximum Projection in the NIS software, exported as TIFF files, and imported into Adobe Photoshop for final presentation.

#### Proximity ligation assay (PLA)

Proximity ligation assay was performed according to the manufacturer’s protocol (Sigma). HCT116:Ki-67-AID were treated with Doxycycline (2μg/ml) and 2 mM thymidine, 24 and 18 h before the addition of Auxin 1000μM, respectively. After the 4 h treatment with Auxin the cells were fixed, permeabilized and blocked with BSA as previously described [60]. The antibodies were used at a concentration as follows, 1:100 anti-MCM3 (clone E-8) (Cat# Sc-390480) and 1:10,000 anti-GFP [PABG1] (Cat# PABG1-20, RRID:AB_2749857). PLA probes were added, and ligation was performed following manufacturer instructions (Sigma). Coverslips were mounted on DAPI and observed on the previously mentioned wide-field NIKON microscope.

#### Proteomics

HCT116 (Ki-67-APEX2) and HCT116 (unmodified) were cultured in SILAC medium (McCoy’s 5A Media for SILAC, Thermo Fisher Scientific); Light (L-Lysine monohydrochloride (Sigma-Aldrich) - L-Arginine monohydrochloride (Sigma-Aldrich) (Ki-67-APEX2) and Heavy (L-Lysine-13C6,15N2 hydrochloride (Sigma-Aldrich) (HCT116, unmodified) respectively for at least 6 passages respectively for at least 6 passages. The cells were treated with 2 mM thymidine for 18 h, and 2.5 mM Biotin added for 30min then the H_2_O_2_ was added for 1min and the reaction was stopped by the addition of Stop Buffer. Cells were then counted and mixed 1:1 (Control;APEX). Cells were lysed for 30min on ice, sonicated and then incubated for 1h at 4°C with streptavidin beads (Pierce). The beads were then washed 3 times with lysis buffer (50mM Tris-HCl, 0.5% IGEPAI, 200mM NaCl) and the biotinylated proteins were eluted with sample buffer and processed for mass spectrometry analysis.

#### Phosphoproteomics

HCT116 (Ki-67-AID) were cultured in SILAC medium (McCoy’s 5A Media for SILAC, Thermo Fisher Scientific); Light (L-Lysine monohydrochloride (Sigma-Aldrich) - L-Arginine monohydrochloride (Sigma-Aldrich) (auxin) and heavy (L-Lysine-13C6,15N2 hydrochloride (Sigma-Aldrich) (control) respectively for at least 6 passages. The cells were treated with Doxycycline (2μg/ml) and 2 mM thymidine, 24 and 18 h before the addition of Auxin 1000μM, respectively. After the 4 h treatment with Auxin the cells were counted and mixed 1:1 (Control/Auxin). Nuclear isolation was performed as described in [61]. Cell lysate was sent for mass spectrometry analysis.

#### Mass spectrometry analyses

For the SILAC APEX experiment (HCT116-APEX vs HCT116-unmodified) proteins were separated on gel (NuPAGE Novex 4-12% Bis-Tris gel, Life Technologies, UK), in NuPAGE buffer (MES) for 10 min and visualised using Instant*Blue*^TM^ stain (AbCam, UK). The stained gel band was excised and de-stained with 50mM ammonium bicarbonate (Sigma Aldrich, UK) and 100% (v/v) acetonitrile (Sigma Aldrich, UK) and proteins were digested with trypsin, as previously described [62]. In brief, proteins were reduced in 10 mM dithiothreitol (Sigma Aldrich, UK) for 30 min at 37°C and alkylated in 55 mM iodoacetamide (Sigma Aldrich, UK) for 20 min at ambient temperature in the dark. They were then digested overnight at 37°C with 13 ng μL-1 trypsin (Pierce, UK). For the SILAC phospho-proteomics experiment, proteins from HCT116 (Ki67-Aid – auxin vs. control) were fully separated on a gel (same as above) and stained under the same conditions. The sample was separated in eight different fractions (i.e. eight gel pieces) and digested under the same conditions described above. Following digestion, samples were diluted with equal volume of 0.1% TFA and spun onto StageTips as described by Rappsilber *et al* (2007) [63]. For the phospho-proteomics experiment, 10% of each fraction was used for the StageTip process and the rest 90% was subjected to a phospho-enrichment process. For those samples prepared for StageTips, peptides were eluted in 40 μL of 80% acetonitrile in 0.1% TFA and concentrated down to 1 μL by vacuum centrifugation (Concentrator 5301, Eppendorf, UK). They were then prepared for LC-MS/MS analysis by diluting it to 5 μL by 0.1% TFA. Phospho-enrichment was done by using the MagReSyn® Ti-IMAC (ReSyn Biosciences, South Africa) according to the manufacturer’s protocol. After the final step of the process, samples were dried under vacuum centrifugation, resuspended in 0.1% TFA and injected as described below.

For the APEX experiment, LC-MS analysis was performed on an Orbitrap Fusion™ Lumos™ (Thermo Fisher Scientific, UK) while for the phospho-proteomics experiment the Orbitrap Exploris™ 480 Mass Spectrometer (Thermo Fisher Scientific, UK) was used, both coupled on-line, to an Ultimate 3000 HPLC (Dionex, Thermo Fisher Scientific, UK). Peptides were separated on a 50 cm (2 µm particle size) EASY-Spray column (Thermo Fisher Scientific, UK), which was assembled on an EASY-Spray source (Thermo Fisher Scientific, UK) and operated constantly at 50°C. Mobile phase A consisted of 0.1% formic acid in LC-MS grade water and mobile phase B consisted of 80% acetonitrile and 0.1% formic acid. Peptides from the APEX experiment and gel fractions were loaded onto the column at a flow rate of 0.3 μL min^-1^ and eluted at a flow rate of 0.25 μL min^-1^ according to the following gradient: 2 to 40% mobile phase B in 120 min and then to 95% in 11 min. Mobile phase B was retained at 95% for 5 min and returned back to 2% a minute after until the end of the run (190 min). For the total proteome samples (8 fractions) the gradient was different; 2 to 40% mobile phase B in 150 min, with a total run time of 190 min.

All MS runs were performed in a Data Dependent Acquisition (DDA) mode. For Orbitrap Fusion™ Lumos™ survey scans were recorded at 120,000 resolution (scan range 350-1500 m/z) with an ion target of 4.0e5, and injection time of 50ms. MS2 was performed in the ion trap at a rapid scan mode, with ion target of 2.0E4 and HCD fragmentation [64] with normalized collision energy of 28. The isolation window in the quadrupole was 1.4 Thomson. Only ions with charge between 2 and 6 were selected for MS2. Dynamic exclusion was set at 60 s. On Orbitrap Exploris™ 480, MS1 scans were recorded at 120,000 resolution (scan range 350-1500 m/z) with an ion target of 3.0e6, and injection time of 50ms. The resolution for the MS2 scans was set at 15,000, with ion target of 8.0E4 and HCD fragmentation with normalized collision energy of 30, and injection time at 64ms. The isolation window in the quadrupole was 1.4 Thomson. Only ions with charge between 2 and 5 were selected for MS2. Dynamic exclusion was set at 60 s.

The MaxQuant software platform [65] version 1.6.1.0 was used to process the raw files and search was conducted against the *Homo sapiens* (released 14/05/19) protein database, using the Andromeda search engine[66]. For the first search, peptide tolerance was set to 20 ppm while for the main search peptide tolerance was set to 4.5 pm. Isotope mass tolerance was 2 ppm and maximum charge to 5. Digestion mode was set to specific with trypsin allowing maximum of two missed cleavages. Carbamidomethylation of cysteine was set as fixed modification. Oxidation of methionine, and phosphorylation of serine, threonine and tyrosine were set as variable modifications. Multiplicity was set to 2, specifying Arg10 and Lys6 for heavy labels. Peptide and protein identifications were filtered to 1% FDR.

#### RNA sequencing

RNA was extracted from HCT116 Ki-67-AID in presence or absence of Ki-67 and extracted using the Monarch Total RNA Miniprep Kit (New England Biolabs, Hitchin, UK) according to manufacturer’s protocol. RNA samples were sent to Macrogen (Japan). Macrogen Europe BV constructed libraries using Illumina TruSeq stranded mRNA library preparation with Ribozero rRNA depletion. Sequencing was performed with a Novaseq 6000 platform, at 50M paired-end reads per sample.

The paired-end raw reads were mapped to the human reference genome GRCh38 using the annotations from GENCODE 28 [67] with HISAT2 under standard conditions. The resulting alignments were filtered for high quality hits using SAMtools v0.1.19 [68] with a minimum selection threshold score of 30. HTSEQ was used to assemble the mapped reads into transcripts and quantify their expression levels.

Deseq2, was used to identify differentially transcribed genes between samples. The differential expression was expressed in the form of log2 fold change between sample and control and deemed statistically significant by a lower p value of 0.05.

Functional enrichment was analysed using String (string-db.org), while Venn diagrams were performed in the open software FunRich. Volcano plots were performed using the ggplot package in R v3.5.0.

#### R2 Genomics analyses

The analysis has been conducted in the R2: Genomics Analysis and Visualization Platform (https://hgserver1.amc.nl). The analysis of the genes has been conducted across data sets in mixed tumors of the u133p2.

### Quantification and statistical analysis

#### FISH analyses

For the intensity analyses, 3D stack images were exported and analysed with ImageJ. ROIs at the size of the nucleus was created, background was subtracted, and the mean intensity was used to generate the violin plots.

For the signal area analyses, 3D stack images were exported and analysed with ImageJ. Scale was set to 1μm, ROIs of the signals were created using threshold function and wand tracing tool, the area in μm2 was used to generate the violin plots.

For the loci distance analyses, 3D stack images were exported and analysed with ImageJ. The in-between distance of the spots was measured and used to generate the violin plots.

For the intensity ratio between the two loci analyses, 3D stack images were exported and analysed with ImageJ. The ROIs at the size of the two signals were created, the ratio of the mean intensity of the two foci was generated and used to generate the violin plots

#### PLA

Spots lying within nuclear masks were counted in control and Ki-67 degradation, the numbers of foci were used to generate the violin plots.

#### Statistical analyses

Statistical analyses were performed either in Excel (Chi-square test) or in R (using the Wilcoxon rank test function, differential expression, lowest smoothing).

## Declarations

### Resource availability

Any additional information required to re-analyse the data reported in this paper is available from the lead contact upon request.

### Lead contact

Further information and requests for resources and reagents should be directed to and will be fulfilled by the lead contact, Paola Vagnarelli.

### Materials availability

All unique/stable reagents generated in this study are available from the lead contact with a completed material transfer agreement.

### Data and code availability

RNA sequencing and proteomic data have been deposited at Arrayepress (Accession E-MTAB-12279) and PXD037513 respectively and are publicly available as of the date of publication. Accession numbers are listed in the key resources table.

This paper does not report original code.

### Author contributions

Conceptualization: PV; Methodology: PV, KS, CS, FH; Formal Analysis: KS, PV, CS; Investigation: KS, PV, CS; Data Curation: KS, CS; Writing – Original Draft: PV, KS; Writing – Review & Editing: PV, KS, CS; Visualization: PV, KS; Supervision: PV, JR; Project Administration: PV; Funding Acquisition: PV.

## Acknowledgments

The authors would like to thank: Prof Masato Kanemaki (Mishima, Shizuoka, Japan) for the HCT116 CMV-OsTIR1 and HCT116 Tet-OsTIR1, Dr Lewis (Brunel University, UK) for the gift of the The HCT116(wt), HCT116 (wt, MLH1 rescued), and SW480 cell lines, Dr Steven D. Cappell (NCI, Bethesda) for the CSII-pEF-hDHB-mCherry plasmid, Prof Eric Schirmer (Edinburgh University, UK) for the IFNβ-firefly plasmid, Prof. Frank van Kuppeveld (Utrecht University, Netherlands) for pCRISPR-hCas-2xSTING-Puro plasmid, Dr Denise Ragusa (Brunel University, UK) for help in selecting the single copy probes, Professor Sala (Brunel University, UK) for useful discussions. The Vagnarelli lab is supported by the Wellcome Trust Investigator award 210742/Z/18/Z to Paola Vagnarelli. KS was supported by a CHMLS PhD scholarship (Brunel University London).

## Figure Legends

**Supplementary Figure 1**

**Generation of AID:mclover endogenously tagged HCT116 cell line.**

A) PCR genotyping of the endogenously tagged AID clones. Gel electrophoresis of the parental cell line (WT) and the targeted cell line (AID) with the indicated primers pairs as shown in (B). M=1kb ladder.

B) Scheme of the expected band sizes obtained by PCR for the wt and targeted alleles with the indicated primers.

C) Western blot of WCL of HCT116:Ki-67-AID untreated treated (-) or treated (+) with Auxin or doxycycline (Dox) for the time indicated. Blots were probed with anti Ki-67 antibody (top) and tubulin (bottom). The lysates were run on an SDS PAGE gradient gel (5-15 %) to resolve all the isoforms of Ki-67. All the isoforms are degraded upon Auxin treatment.

D) Top: Scheme of the experiment. EdU was added 30 minutes before each time point. Bottom: The graph represents the percentage of EdU positive cells at the different time points. The values are the mean of 3 biological replicas and the error bars represent the standard deviations. Sample size: Control 3h=1060, 4h=1038, 6h=994; Auxin 3h=3572, =3503, 6h=3444 The data were statistically analysed with a Chi-squared test (control vs auxin). ***= p<0.001

E) Top: Scheme of the experiment. Bottom: Distribution of the replicating cells according to the patterns shown in Figure 1 G. The values are the mean of 2 biological replicas and the error bars represent the standard deviations. Sample size: Control=270, Oligo 1#=561, Oligo 2#=291. The data were statistically analysed with a Chi-squared test (control vs auxin). ***= p<0.001

F) Top: Scheme of the experiment. Thy = Thymidine, Bottom: Representative images of Nucleolin immunostaining using anti Nucleolin antibodies on HCT116:Ki-67-AID cell line without (top) of with (bottom) Auxin. Scale bar 5 μm.

**Supplementary Figure 2**

**Ki-67 degradation alters gene expression when cells exit mitosis in its absence.** A) Volcano plot of the differentially expressed genes when Ki-67 is degraded in cells arrested in nocodazole and then released in Thymidine with or without Auxin for 18 h. The pink line represents p-value < 10e^-20^, the Hot pink line p-value=10e^-20^-10e^-30^ and the purple line p-value=10e^-20^-10e^-60^. Several genes are differentially expressed in this sample. B) Table of DNA replication genes downregulated in G1 upon Ki-67 degradation in mitosis. C) Venn diagram comparing the numbers of upregulated and downregulated genes in the RNA-seq experiment in (A) (Mitosis-G1) and in Figure 3 A (G1). D) Magic analyses of upregulated genes using a matrix containing the gene bodies (from the promoter to the end of the last exon and 1kb flanking sequence either side of the gene body). E-I) Western blots of whole cell lysate of the indicated cell lines upon Control or Ki-67 RNAi. Oligo #1 and Oligo #2 indicate two already published SiRNA oligos. The blots were probed with anti Ki-67 (top panels) and anti alpha tubulin (bottom panels) antibodies.The graphs show the quantification of the blots.

**Supplementary Figure 3**

A) PCR genotyping of the endogenously tagged APEX2 clones. Gel electrophoresis of the parental cell line (WT) and the targeted cell line (APEX2) with the indicated primers pairs as shown in (B). M=1kb ladder

B) Scheme of the expected band size obtained by PCR for the wt and targeted alleles with the indicated primers.

C) Line plots (right) of the nucleus of HCT116:Ki-67-AID across the yellow line show in the images (left panels). Scale bar 5 μm.

D) Line plots of Ki-67 foci and BrdU foci across the selected regions in (4 K). Scale bar 5 μm.

**Supplementary Figure 4**

**Ki-67 degradation at G1/S leads to CDK2 inactivation.**

A) Scheme of the experiment in (B). Thy = Thymidine B) (Top) Representative Western blot of HCT116: Ki-67-AID cell line blocked with Thymidine then untreated (Ki-67 AID) or treated with Auxin for 4h (Ki-67 AID Auxin) in the presence (+) or absence (-) of BI8626. The blots were probed with anti GAPDH and an tiChk1 (B) antibodies. The graphs at the top represent the quantification of the blots. The values represent the average of 3 independent replicas and the error bars are the standard deviations. The experiments were analysed by a Student’s t-test. *=p<0.05, ***= p< 0.001.

C) Scheme of the experiment (top) and representative images of the DHB-mCherry reporter localisation in cells at the G1/S boundary with or without Ki-67 (bottom). Thy = Thymidine. Scale bar 10 μm.

D) Quantification of the experiment in (C). The violin plots represent the distribution of the cytoplasmic/nuclear mean signals of 3 biological replicates. The box inside the violin represents the 75^th^ and 25^th^ percentile, whiskers are the upper and lower adjacent values and the line is the median. N: Control=336 Auxin=322. The data were analysed with a Wilcoxon test. ***=p<0.001.

**Supplementary Figure 5**

**Correlation between Ki-67 and the interferon pathway.**

A) Scheme of the expected band size obtained by PCR for the wt and targeted alleles with the indicated primers.

B) PCR genotyping of the Ki-67-AID-STING-KO cell line. The black and blue arrows indicate the wt and the KO alleles respectively (as shown in A). M =kb ladder.

C-D) R2 genomics analyses of patient and experimental data sets for IFIT1 (C) and STAT2 (D) compared against MKI67 gene expression, X and Y-axis the log2 transformed average expression level and the standard deviation is represented.

## Supplemental Method

**Supplementary Table 1:**
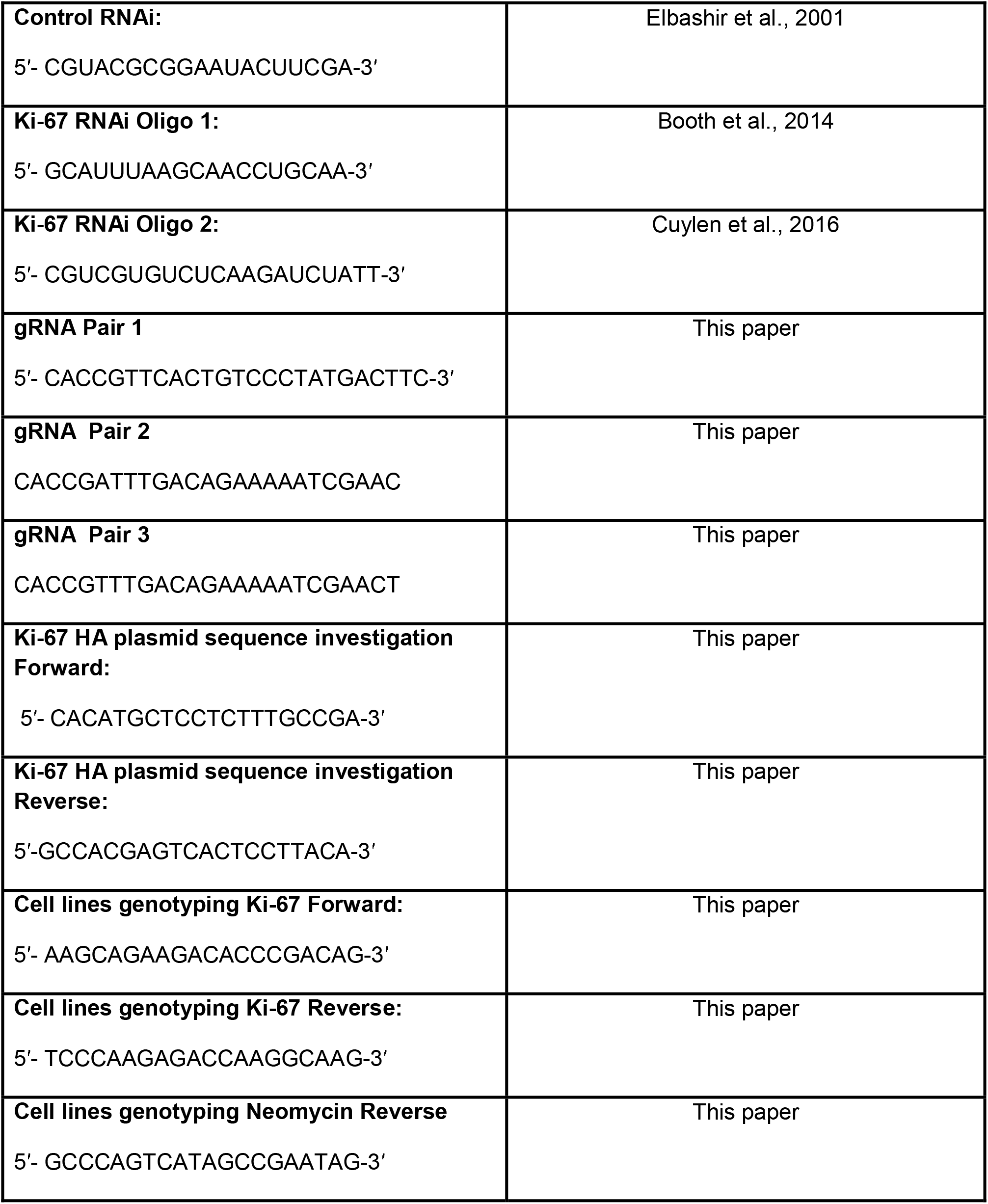
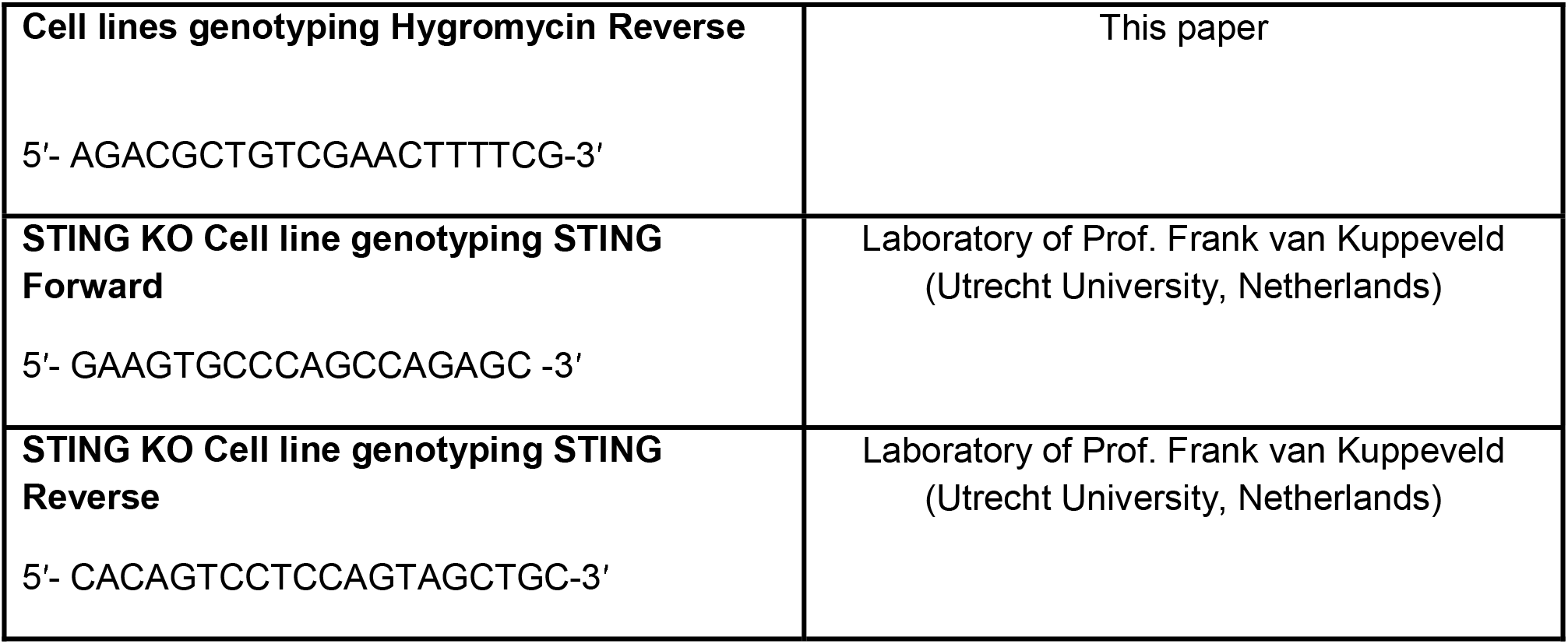
Oligonucleotides used in this study

**Supplementary Table 2:** List of Antibodies, reagents and cell lines in this study

| REAGENT or RESOURCE | SOURCE | IDENTIFIER |
| --- | --- | --- |
| **Antibodies** |  |  |
| Mouse monoclonal anti-MCM3(clone E-8) | Santa Cruz Biotechnology | Cat# Sc-390480 |
| Mouse monoclonal anti-PCNA (clone PC10) | Santa Cruz Biotechnology | Cat# Sc-56<br>RRID:AB_628110 |
| Mouse monoclonal anti-ORC1 (clone F-10) | Santa Cruz Biotechnology | Cat# Sc-398734 |
| Mouse monoclonal anti-p21 Waf1:Cip1 (clone F-5) | Santa Cruz Biotechnology | Cat# Sc-6246<br>RRID:AB_628073 |
| Mouse monoclonal anti-p53 (clone DO-1) | SIGMA | Gift from Dr. Evgeny Makarov |
| Mouse monoclonal anti-c-Myc (clone C-33) | Santa Cruz Biotechnology | Cat#Sc-42<br>RRID:AB_2282408 |
| Mouse monoclonal anti-a-tubulin (clone B-5-1-2) | Sigma-Aldrich | Cat# T5168<br>RRID:AB_477579 |
| Mouse monoclonal anti-GAPDH | Proteintech | Cat# 60004-1-Ig,<br>RRID:AB_2107436 |
| Rabbit polyclonal anti GFP [PABG1] | Chromotek | Cat# PABG1-20,<br>RRID:AB_2749857 |
| Mouse monoclonal anti-BrdU Alexa Fluor® 647 (clone 3d4) | Biolegent | Cat# 364108<br>RRID:AB_2566452 |
| Mouse monoclonal anti-Ki-67 | BD Biosciences | Cat# 610968<br>RRID:AB_398281 |
| Rabbit polyclonal anti-Nucleolin | Abcam | Cat# 22758<br>RRID:AB_776878 |
| Rabbit monoclonal anti-Histone H2A.X (clone D17A3) | Cell signalling | Cat# 7631<br>RRID:AB_10860771 |
| Mouse monoclonal anti-p-Histone H2A.X Antibody (Ser 139) | Santa Cruz Biotechnology | Cat# Sc-517348<br>RRID:AB_2783871 |
| Rabbit polyclonal anti-Histone H3 | Active Motif | Cat# 39163<br>RRID:AB_2614978 |
| Mouse monoclonal anti-H3K27me2/3 | Active Motif | Cat# 39535<br>RRID:AB_2793246 |
| Rabbit polyclonal anti-H3K9me3 | Active Motif | Cat# 39161<br>RRID:AB_2532132 |
| Mouse monoclonal anti-H4K20me1 | Active Motif | Cat# 39727<br>RRID:AB_2615074 |
| Rabbit polyclonal anti-H3K9me2 | Active Motif | Cat# 39239<br>RRID:AB_2793199 |
| Mouse polyclonal Texas Red | Jackson Immunoresearch | Cat# 715-585-150<br>RRID:AB_2340854 |
| Goat polyclonal anti-Mouse HRP | Thermo Fisher Scientific | Cat# 31444<br>RRID:AB_228321 |
| Goat polyclonal anti-Mouse 800CW | LI-COR | Cat# 926-32210<br>RRID:AB_621842 |
| Goat polyclonal anti-Rabbit 680RD | LI-COR | Cat# 926-68071<br>RRID:AB_10956166 |
| **Chemicals, peptides, and recombinant proteins** |  |  |
| 3-Indoleacetic acid | Sigma-Aldrich | Cat# I2886<br>CAS:87-51-4 |
| Thymidine | Sigma-Aldrich | Cat# T9250<br>CAS:50-89-5 |
| RO-3306 | Sigma-Aldrich | Cat# SML0569<br>CAS:872573-93-8 |
| Nocodazole | Sigma-Aldrich | Cat# 31430-18-9<br>CAS:87-51-4 |
| MG132 | Millipore | Cat# 474787<br>CAS:133407-82-6 |
| Doxycycline | Sigma | Cat# D9891<br>CAS: 24390-14-5 |
| Palbociclib (PD-0332991) | Selleckchem | Cat# S1116<br>CAS: 827022-32-2 |
| HUWE1 inhibitor BI8626 | MedChemExpress | Cat# HY-120204<br>CAS: 1875036-75-1 |
| **Critical commercial assays** |  |  |
| Click-iT™ EdU Cell Proliferation Kit for Imaging, Alexa Fluor™ 647 | Thermo Fisher Scientific | Cat# C10340 |
| Dual-Luciferase Reporter Kit | Promega | Cat# E1910 |
| Duolink® In Situ PLA® Probe Anti-Rabbit MINUS | Sigma-Aldrich | Cat# DUO92005 |
| Duolink® In Situ PLA® Probe Anti-Mouse PLUS | Sigma-Aldrich | Cat# DUO92001 |
| Duolink® In Situ Detection Reagents Red | Sigma-Aldrich | Cat# DUO92008 |
| Monarch Total RNA Miniprep Kit | New England BioLabs | Cat# T2010S |
| JetPrime® | Polyplus transfection® | Cat# 101000015 |
| Nick Translation Kit | Abbott | Cat# 07J00-001 |
| pGEM-T Easy empty vector | Promega | Cat# A1360 |
| **Deposited data** |  |  |
| RNA sequencing data | This paper | E-MTAB-12279 |
| Proteomics data | This paper | PXD037513 |
| Human reference genome GRCh38 | Genome Reference Consortium Human Build 38 | https://www.ncbi.nlm.nih.gov/assembly/GCF_000001405.26/ |

| **Experimental models: Cell lines** |  |  |
| --- | --- | --- |
| HCT116(wt) | Laboratory of Dr Anabelle Lewis (Brunel University of London, UK) | ATCC, CCL-247 |
| HCT116 (wt, MLH1 rescued) | Laboratory of Dr Anabelle Lewis (Brunel University of London, UK) | Horizon Discovery |
| SW480 | Laboratory of Dr Anabelle Lewis (Brunel University of London, UK) | ATCC, CCL-228 |
| hTERT-RPE1 | Laboratory of Dr Viji Draviam (Queen Mary University of London, UK) | ATCC, CRL-4000 |
| HCT116 OsTR1 | Laboratory of Dr Masato T. Kanemaki (University of Tokyo, Japan) | N/A |
| HCT116:Ki-67-AID | This paper | N/A |
| HCT116:Ki-67-AID mCherry-DHB | This paper | N/A |
| HCT116:Ki-67-APEX2 | This paper | N/A |
| HCT116:Ki-67-AID STING-KO | This paper | N/A |
| HeLa Kyoto | ATCC | CRL-11268 |
| **Oligonucleotides** |  |  |
| See Table S1 for Oligonucleotides used in this study |  |  |
| **Recombinant DNA** |  |  |
| pX330-U6-Chimeric_BB-CBh-hSpCas9 | [55] | Addgene plasmid #42230 |
| pMK289 (mAID-mClover-NeoR) | [48] | Addgene plasmid #72827 |
| pMK290 (mAID-mClover-Hygro) | [48] | Addgene plasmid #72828 |
| APEX2-csChBP | [69] | Addgene plasmid #108876 |
| pBluescript II SK(+)-Ki-67 | This paper | Biomatik |
| pBluescript II SK(+)-Ki-67 - mAID-mClover-Neo | This paper | N/A |
| pBluescript II SK(+)-Ki-67 - mAID-mClover-Hygro. | This paper | N/A |
| pX330-U6-Chimeric-BB-CBh-hSpCas9-gRNA-P1 | This paper | N/A |
| pX330-U6-Chimeric-BB-CBh-hSpCas9-gRNA-P2 | This paper | N/A |
| pX330-U6-Chimeric-BB-CBh-hSpCas9-gRNA-P3 | This paper | N/A |
| pBluescript II SK(+)-Ki-67-APEX2-mClover-Neo | This paper | N/A |
| pBluescript II SK(+)-Ki-67-APEX2-mClover-Hygro | This paper | N/A |
| pCRISPR-hCas-2xSTING-Puro | Laboratory of Prof. Frank van Kuppeveld (Utrecht University, Netherlands) | N/A |
| pMSCV-Blasticidin | [70] | Adgene plasmid #75085 |
| CSII-pEF-hDHB-mCherry | Laboratory of Dr<br>Steven D. Cappell<br>(Center for Cancer<br>Research<br>National Cancer<br>Institute, USA) | N/A |
| Software and algorithms |  |  |
| ImageJ | [71] | https://imagej.nih.gov/ij/ |
| Samtools | [68] | http://samtools.sourceforge.net/ |
| HTSeq | [72] | https://htseq.readthedocs.io/en/master/ |
| Deseq2 | [73] | https://bioconductor.org/packages/release/bioc/html/DESeq2.html |
| NIS-Elements | Nikon |  |
| SoftWoRx | Delta Vision |  |
| R studio |  | https://posit.co/ |
| MaxQuant software platform | [65] |  |
| Andromeda search engine | [66] |  |

