## Supplementary figures and images for "Ki-67 is necessary during DNA replication for forks protection and genome stability"

### Supplementary Figure 1

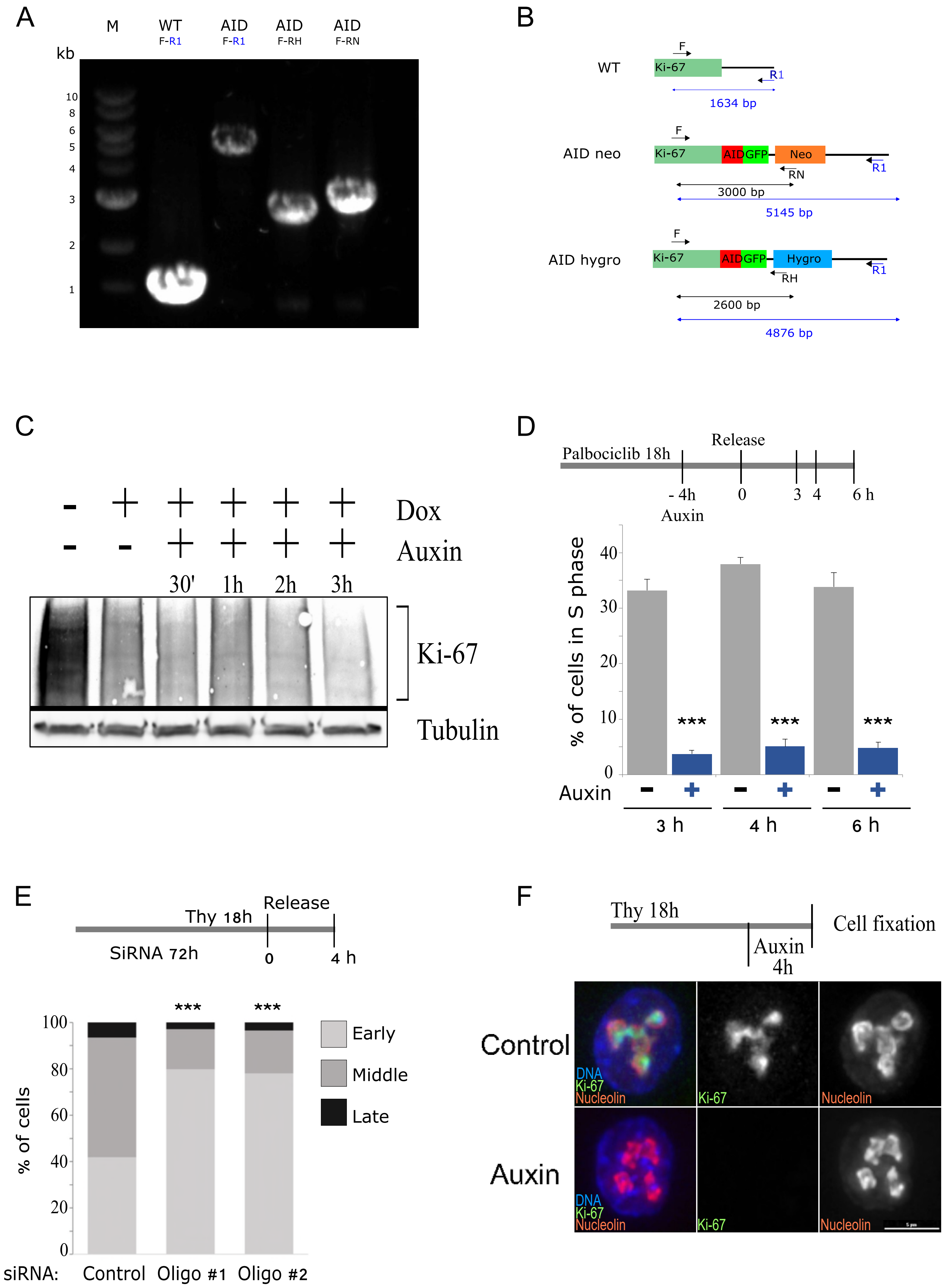

### Supplementary Figure 2

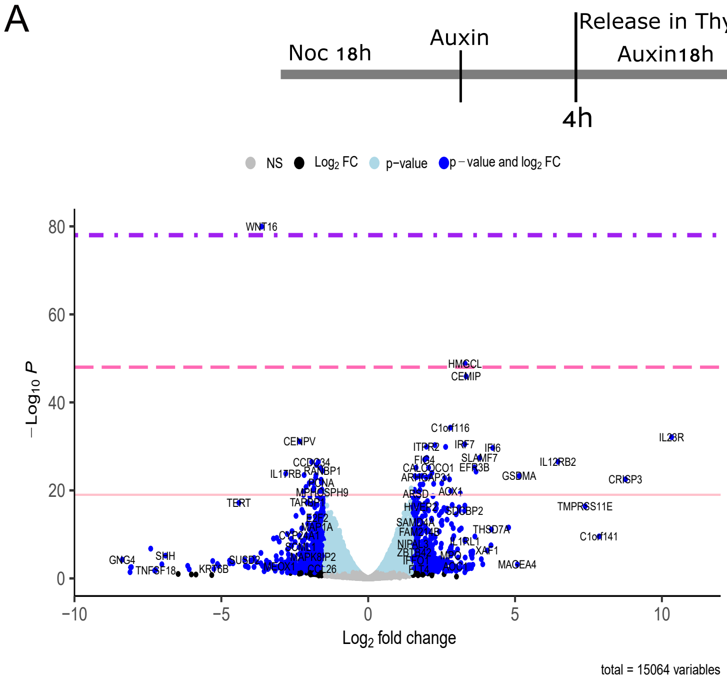


B

| | log2 Fold Change |
|---|---|
| **PCNA** | -1.592506284 |
| **MCM7** | -1.602911138 |
| **ORC6** | -1.735630858 |
| **CDC45** | -1.750303313 |
| **ORC1** | -1.817978697 |

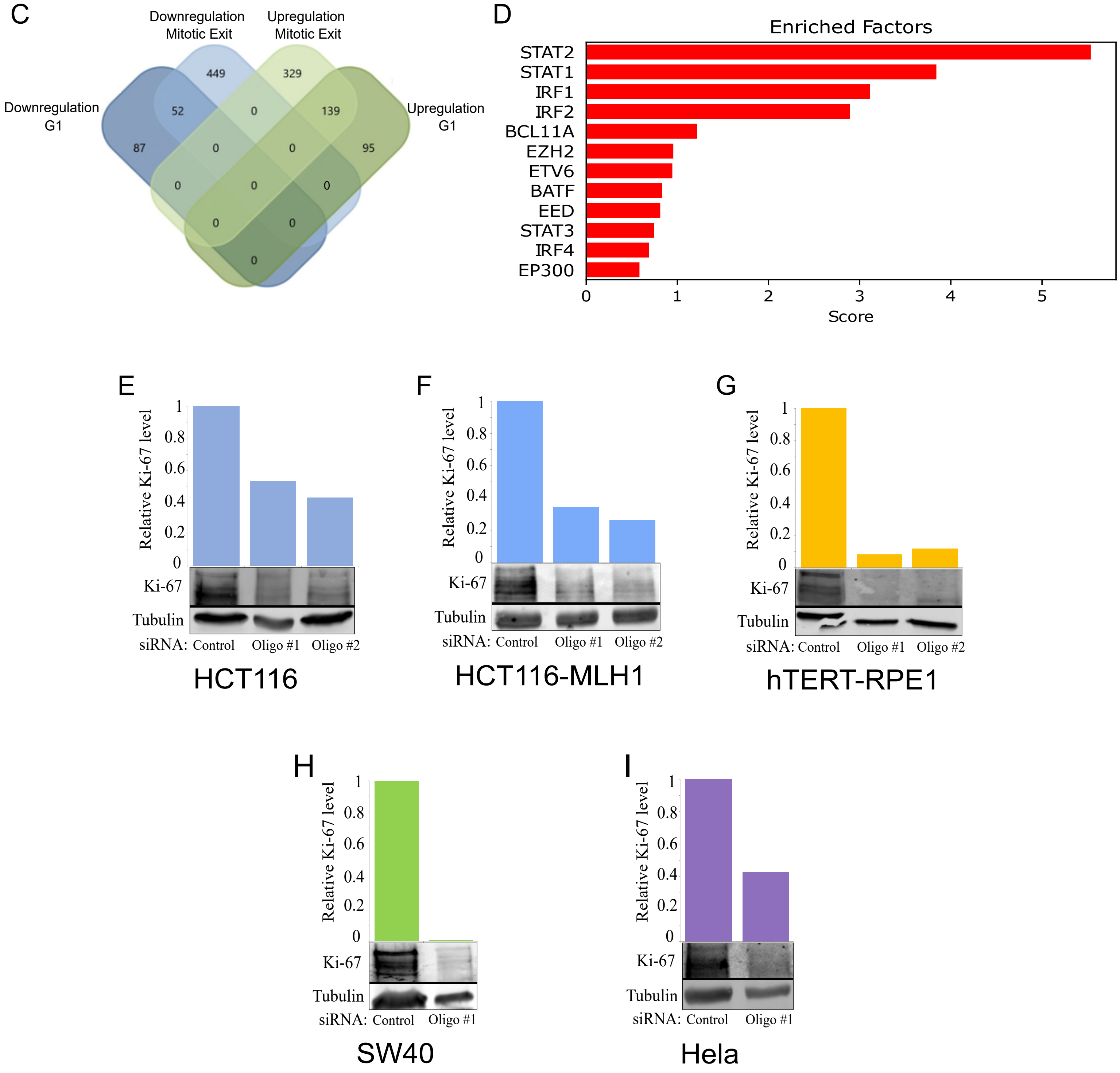

### Supplementary Figure 3

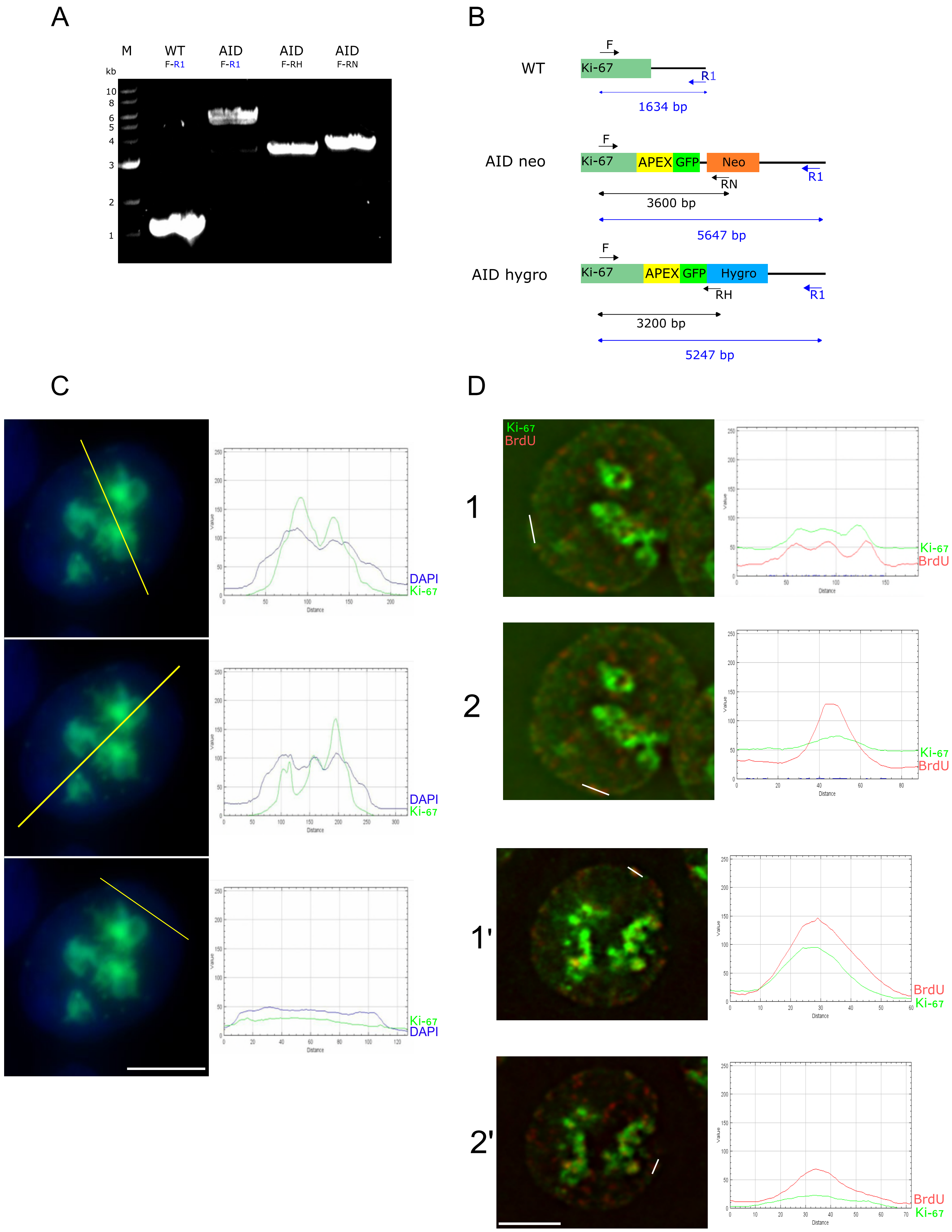

### Supplementary Figure 4

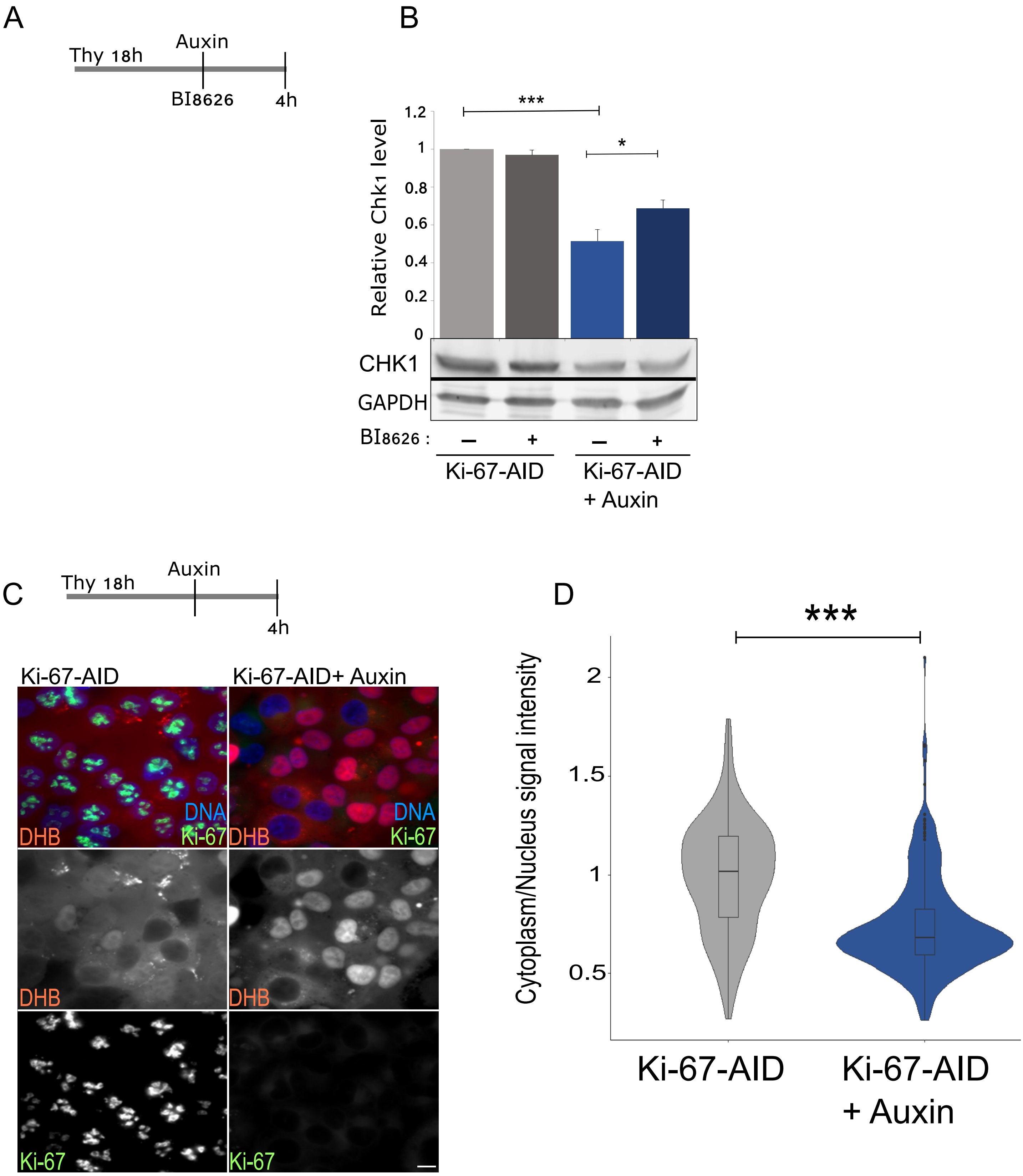

### Supplementary Figure 5

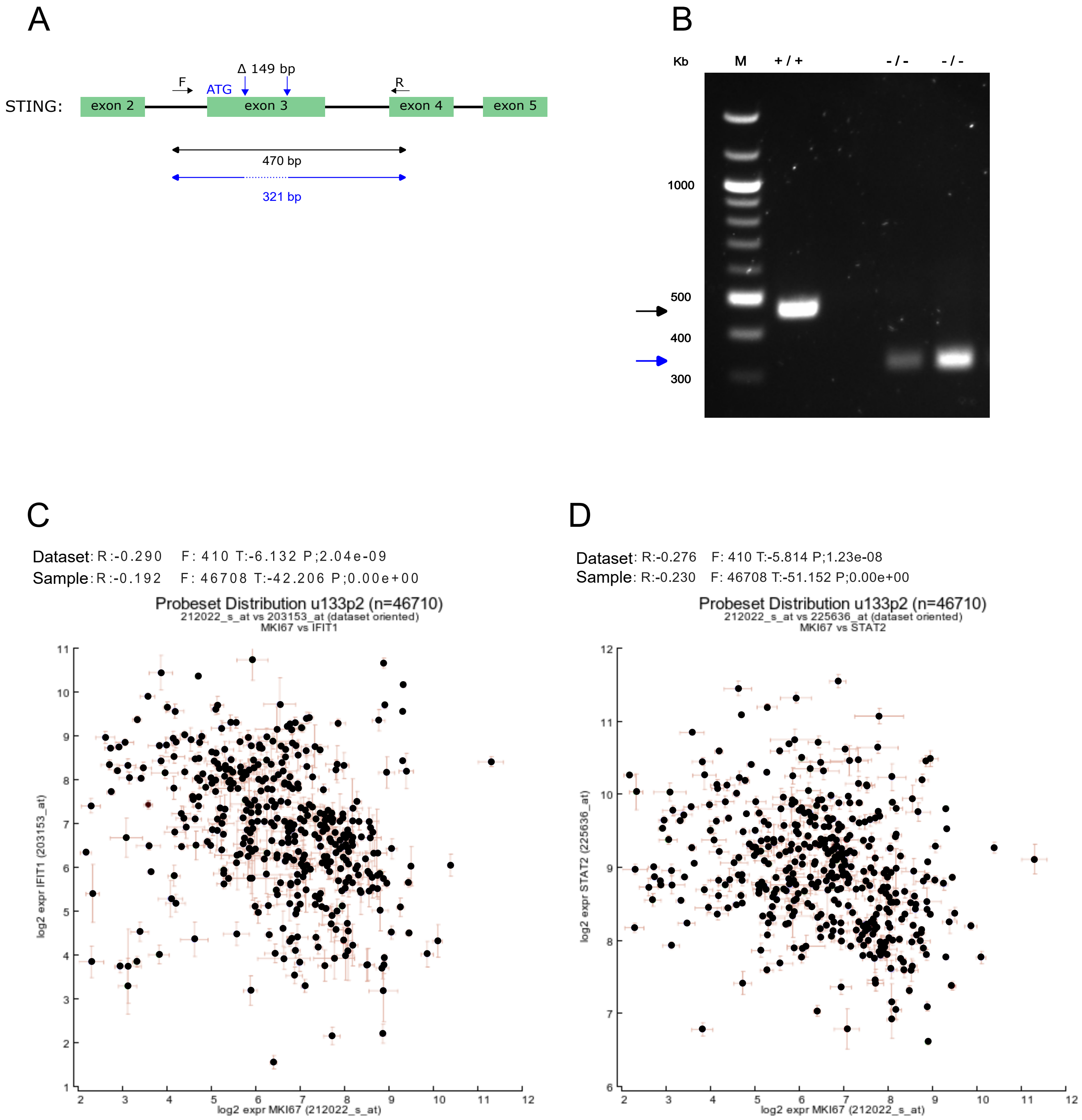
